# PLETHORA-WOX5 interaction and subnuclear localisation control *Arabidopsis* root stem cell maintenance

**DOI:** 10.1101/818187

**Authors:** Rebecca C. Burkart, Vivien I. Strotmann, Gwendolyn K. Kirschner, Abdullah Akinci, Laura Czempik, Anika Dolata, Alexis Maizel, Stefanie Weidtkamp-Peters, Yvonne Stahl

## Abstract

Maintenance and homeostasis of the stem cell niche (SCN) in the *Arabidopsis* root is essential for growth and development of all root cell types. The SCN is organized around a quiescent center (QC) maintaining the stemness of cells in direct contact. The key transcription factors (TFs) WUSCHEL-RELATED HOMEOBOX 5 (WOX5) and PLETHORAs (PLTs) are expressed in the SCN where they maintain the QC and regulate distal columella stem cell (CSC) fate. Here, we describe the concerted mutual regulation of the key TFs WOX5 and PLTs on a transcriptional and protein interaction level. Additionally, by applying a novel SCN staining method, we demonstrate that both WOX5 and PLTs regulate root SCN homeostasis as they control QC quiescence and CSC fate interdependently. Moreover, we uncover that some PLTs, especially PLT3, contain intrinsically disordered prion-like domains (PrDs) that are necessary for complex formation with WOX5 and its recruitment to subnuclear microdomains/nuclear bodies (NBs) in the CSCs. We propose that this partitioning of PLT-WOX5 complexes to NBs, possibly by phase separation, is important for CSC fate determination.

## Introduction

The root system of higher plants is essential for plant life, as it provides anchorage in the soil and access to nutrients and water. It arises from a population of long-lasting stem cells residing in a structure called root apical meristem (RAM) at the tip of the root. Within the *Arabidopsis thaliana* RAM, the stem cell niche (SCN) consists of on average four to eight slowly dividing cells, the QC cells, which act as a long-term reservoir and signalling center by maintaining the surrounding shorter-lived, proliferating stem cells (also called initials) in a non-cell autonomous manner (van den Berg et al, 1997; Lu et al, 2021). These stem cells continuously divide asymmetrically, thereby generating new stem cells that are still in contact with the QC. The hereby-produced daughter cells frequently undergo cell divisions and are shifted further away from the QC to finally differentiate into distinct cell fates. By this mechanism, the position of the stem cells in the root remains the same throughout development and their precise orientation of division leads to the formation of concentrically organized clonal cell lineages representing a spatio-temporal developmental gradient (Dolan et al, 1993; van den Berg et al, 1997; Benfey & Scheres, 2000). From the inside to the outside the following root cell tissues develop: vasculature, pericycle, endodermis, cortex and epidermis plus columella and lateral root cap at the distal root tip (Fig. 1A).

**Figure 1 -.**
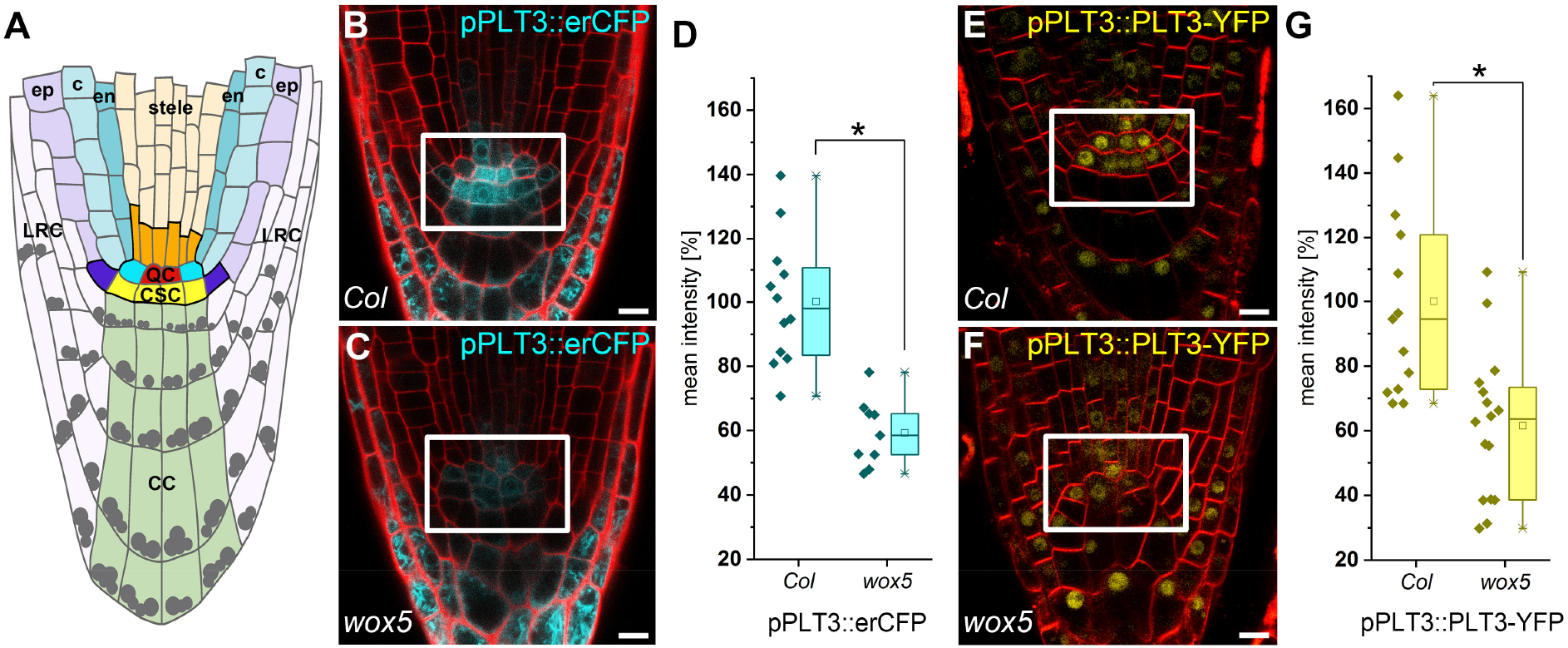
WOX5 positively regulates PLT3 expression. **A**, Schematic representation of the *Arabidopsis* root meristem. The QC cells (red) maintain the surrounding stem cells (initials) outlined in black together building the root stem cell niche (SCN). The different cell types are colour-coded. QC = quiescent center (red); CSC = columella stem cells (yellow); CC = columella cells (green); LRC = lateral root cap (light purple); ep = epidermis (purple); c = cortex (light turquoise); en = endodermis (dark turquoise); bright turquoise = cortex/endodermis initials; dark purple = epidermis/lateral root cap initals; dark orange = stele initials; stele = light orange; grey dots = starch granules. **B**, **C**, Representative images of pPLT3::erCFP (cyan) expressing and PI-stained (red) *Arabidopsis* roots in *Col* or *wox5* background, respectively. **D**, Mean fluorescence intensities of the pPLT3::erCFP roots summarized in box and scatter plots. The mean fluorescence intensity of the CFP signal in *Col* roots was to set to 100 %. **E**, **F**, Representative images of pPLT3::PLT3-YFP (yellow) expressing and FM4-64-stained (red) *Arabidopsis* roots in *Col* or *wox5* background, respectively. **g**, Mean fluorescence intensities of the pPLT3::PLT3-YFP expressing roots summarized in box and scatter plots. The mean fluorescence intensity of the YFP signal in *Col* roots was to set to 100%. **D**, **G**, Box = 25-75 % of percentile, whisker = 1.5 interquartile range, − = median, □ = mean value, × = minimum/maximum. After confirming normal distribution of the data by Kolmogorov-Smirnov testing, the data was statistically analyzed by one-way ANOVA and Holm-Sidak post-hoc multiple comparisons test. Asterisks indicate statistically significant differences (α = 0.01), number of analyzed roots n = 9-16. **B**, **C**, **E**, **F**, Scale bars represent 10 μm. SCN = stem cell niche; PI = propidium iodide; YFP = yellow fluorescent protein; CFP = cyan fluorescent protein.

The necessary longevity and continuous activity of the RAM can only be achieved if its stem cell pool is constantly replenished, since cells are frequently leaving the meristematic region due to continuous cell divisions. Therefore, complex regulatory mechanisms involving phytohormones and key TFs regulate stem cell maintenance and the necessary supply of differentiating descendants (Drisch & Stahl, 2015). Here, the APETALA2-type PLT TF family and the homeodomain TF WOX5 play important roles (Aida et al, 2004; Sarkar et al, 2007). WOX5 is expressed mainly in the QC, but maintains the surrounding stem cells non-cell-autonomously by repressing their differentiation (Sarkar et al, 2007; Pi et al, 2015). Loss of WOX5 causes the differentiation of the distal CSCs into starch-accumulating columella cells (CCs), while increased WOX5 expression causes CSC over-proliferation. Hence, WOX5 abundance is critical and necessary to suppress premature CSC differentiation (Sarkar et al, 2007; Pi et al, 2015). WOX5 also represses QC divisions, maintaining the quiescence of the QC by repressing CYCLIN D (CYCD) activity within the QC (Forzani et al, 2014).

The auxin-induced PLTs form a clade of six TFs, and act as master regulators of root development, as multiple *plt* mutants fail to develop functional RAMs (Aida et al, 2004; Galinha et al, 2007; Mähönen et al, 2014). PLT1, 2, 3 and 4 are expressed mainly in and around the QC and form an instructive gradient, which is required for maintaining the balance of stem cell fate and differentiation. This PLT gradient is also necessary for separating auxin responses in the SCN, for the correct positioning of the QC, and the expression of QC markers (Aida et al, 2004; Galinha et al, 2007; Mähönen et al, 2014). Genetically, WOX5 and PLT1 were shown to play an interconnected role in auxin-regulated CSC fate, whereas PLT1 and PLT3 were found to directly positively regulate WOX5 expression (Ding & Friml, 2010; Shimotohno et al, 2018).

Although PLTs and WOX5 are known for controlling stem cell regulation and maintenance in the *Arabidopsis* RAM and genetic evidence for cross regulation exists, the underlying molecular mechanisms are until now largely elusive. Here, we show for the first time that the mutual regulation of expression, but importantly also the ability of PLTs to directly interact with and recruit WOX5 to NBs in CSCs controls stem cell homeostasis in the *Arabidopsis* RAM. NBs are membrane-less, self-assembling protein/RNA containing compartments thought to enhance a variety of physiological reactions upon differential cues (Meyer, 2020; Mao et al, 2011). Therefore, we propose a model in which differential PLT/WOX5 complexes depending on their subnuclear localisation in NBs or in the nucleoplasm regulate stem cell fate in the RAM, possibly by phase separation.

## Results

WOX5 and PLTs are essential players in distal stem cell maintenance (Aida et al, 2004; Pi et al, 2015; Sarkar et al, 2007; Galinha et al, 2007). This, as well as their overlapping expression and protein localisation domains in the root SCN raised the question if they could act together in distal stem cell regulation, where, in comparison to all the other PLTs, particularly PLT3 is highly expressed (Fig. 1B) (Galinha et al, 2007). First, we tested if WOX5 influences *PLT3-*expression. Both a transcriptional and translational PLT3 fluorescent reporter line showed a reduced expression in the QC and CSC in a *wox5* mutant background to around 60 % compared to the *Col-0* (*Col*) wild type roots (Fig. 1B-G, Suppl. Table 5). This extends the previously reported regulation of *PLT1* expression by WOX5 (Ding & Friml, 2010) and shows that WOX5 positively regulates expression of several *PLTs*. To test if *WOX5* expression also depends on PLTs, we produced a transcriptional reporter, which expresses a nuclear-localised mVenus under control of the *WOX5* promoter. In agreement with previous reports, expression of *WOX5* in our transcriptional reporter line is confined to the QC and is only weakly expressed in the stele initials (Sarkar et al, 2007; Pi et al, 2015) (Fig. 2A).

**Figure 2 -.**
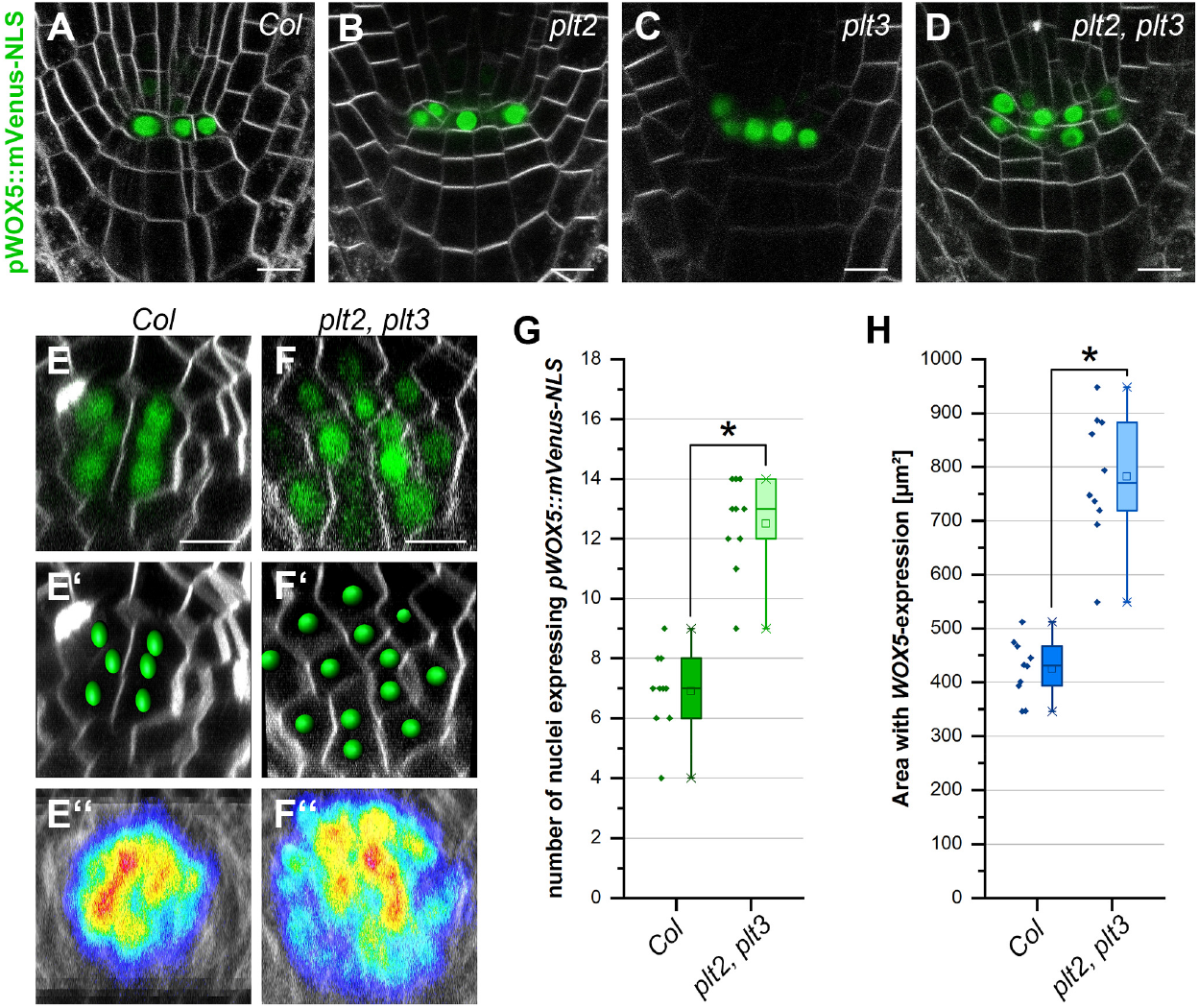
PLTs constrain the WOX5 expression domain. **A**-**F**, Representative FM4-64-stained *Arabidopsis* roots (grey) expressing pWOX5::mVenus-NLS (green) in *Col*, *plt2*, *plt3* and *plt2*, *plt3* double mutant background in longitudinal (**A**-**D)**, or transversal (**E**-**F**) optical sections. **E’**, **F’**, Analysis of representative images in **(E**) and (**F**) in Imaris to detect and count individual expressing nuclei. **E’’**, **F’’**, Overlay of 10 roots showing the area of detected fluorescence (high levels in red, low levels in blue) in *Col* and *plt2*, *plt3* double mutant roots. **G**, Number of nuclei expressing pWOX5::mVenus-NLS in *Col* and *plt2*, *plt3* double mutant roots summarized in box and scatter plots. **H**, Area of WOX5 expression in μm^2^ in *Col* and *plt2, plt3* double mutant roots summarized in box and scatter plots. **G, H** Box = 25-75 % of percentile, whisker = 1.5 interquartile range, − = median, □ = mean value, × = minimum/maximum. After confirming normal distribution of the data by Kolmogorov-Smirnov testing, the data was statistically analyzed by one-way ANOVA and post-hoc Holm-Sidak multiple comparisons test. Asterisks indicate statistically significant differences (α = 0.01). Number of analysed roots n = 10. Scale bars represent 10 μm; NLS = nuclear localisation signal.

In *plt2* and *plt3* single mutants, we observed additional mVenus-expressing cells in the QC region, which may derive from aberrant periclinal cell divisions of the QC (Fig. 2B, C, Suppl. Table 6). This effect is even stronger in the *plt2*, *plt3* double mutant roots, where extra cells are found in all observed roots and often even form an additional cell layer of *WOX5* expressing cells (Fig. 2D). Previously, it was reported that the *Arabidopsis* wild type QC is composed of four to eight cells with a low division rate (Lu et al, 2021; Truernit et al, 2008; Cruz-Ramírez et al, 2013; Stahl et al, 2013). We quantified the number of *WOX5* expressing cells and the area of *WOX5* expression per root by acquiring transverse optical sections through the roots. We observed four to nine *WOX5* expressing cells in the *Col* wild type (Fig. 2E, G, Suppl. Table 6), whereas we found nine to 14 *WOX5* expressing cells and a laterally expanded *WOX5* expression domain in the *plt2*, *plt3* double mutants (Fig. 2F, G, H, Suppl. Table 6). Taken together, our data show that WOX5 positively regulates *PLT* expression, here shown for PLT3, whereas PLT2 and PLT3 redundantly restrict *WOX5* expression to a limited number of cells at QC position, possibly by negative feedback regulation. These observations are in agreement with a previous report, where a role for PLT1 and PLT2 in confining *WOX5* expression was reported (Sarkar et al, 2007).

QC cells rarely divide as they provide a long-term reservoir to maintain the surrounding stem cells (Cruz-Ramírez et al, 2013; Vilarrasa-Blasi et al, 2014). As WOX5 and PLTs control QC cell divisions and CSC maintenance (Sarkar et al, 2007; Forzani et al, 2014; Pi et al, 2015; Aida et al, 2004; Galinha et al, 2007; Mähönen et al, 2014), we asked if these two aspects are interdependent. Therefore, we analysed the cell division rates in the QC and the CSC phenotypes in wild type and mutant roots. To assess these two phenotypes and to probe for their interdependency, we needed to measure the number of dividing QC cells and CSC layers within the same root simultaneously. To enable this, we established a novel staining method, named SCN staining, by combining the 5-ethynyl-2’-deoxyuridine (EdU) and modified pseudo Schiff base propidium iodide (mPS-PI) stainings to simultaneously visualise cell divisions, starch granule distribution as well as cell walls within the same root (Truernit et al, 2008; Cruz-Ramírez et al, 2013). Applying this new staining combination, potential correlations between QC-divisions and CSC cell fates can be uncovered. The EdU-staining is used to analyse QC-divisions by staining nuclei that have gone through the S-phase, detecting cells directly before, during and after cell division (Cruz-Ramírez et al, 2013). However, cell layers and different cell types are hard to distinguish using only EdU staining due to the lack of cell wall staining. Therefore, we additionally applied the mPS-PI-method to stain cell walls and starch which is commonly used for CC and CSC cell fate determination (Truernit et al, 2008; Stahl et al, 2013; Stahl et al, 2009). CCs are differentiated, starch granule-containing cells in the distal part of the root mediating gravity perception. They derive from the CSCs that form one or, directly after cell division, two cell layers distal to the QC. The CSCs lack big starch granules and can thereby easily be distinguished from the differentiated CCs by mPS-PI staining (Truernit et al, 2008; Stahl et al, 2009; Stahl et al, 2013) (see Fig. 3A, B, I, raw data see Suppl. Table 11).

**Figure 3 -.**
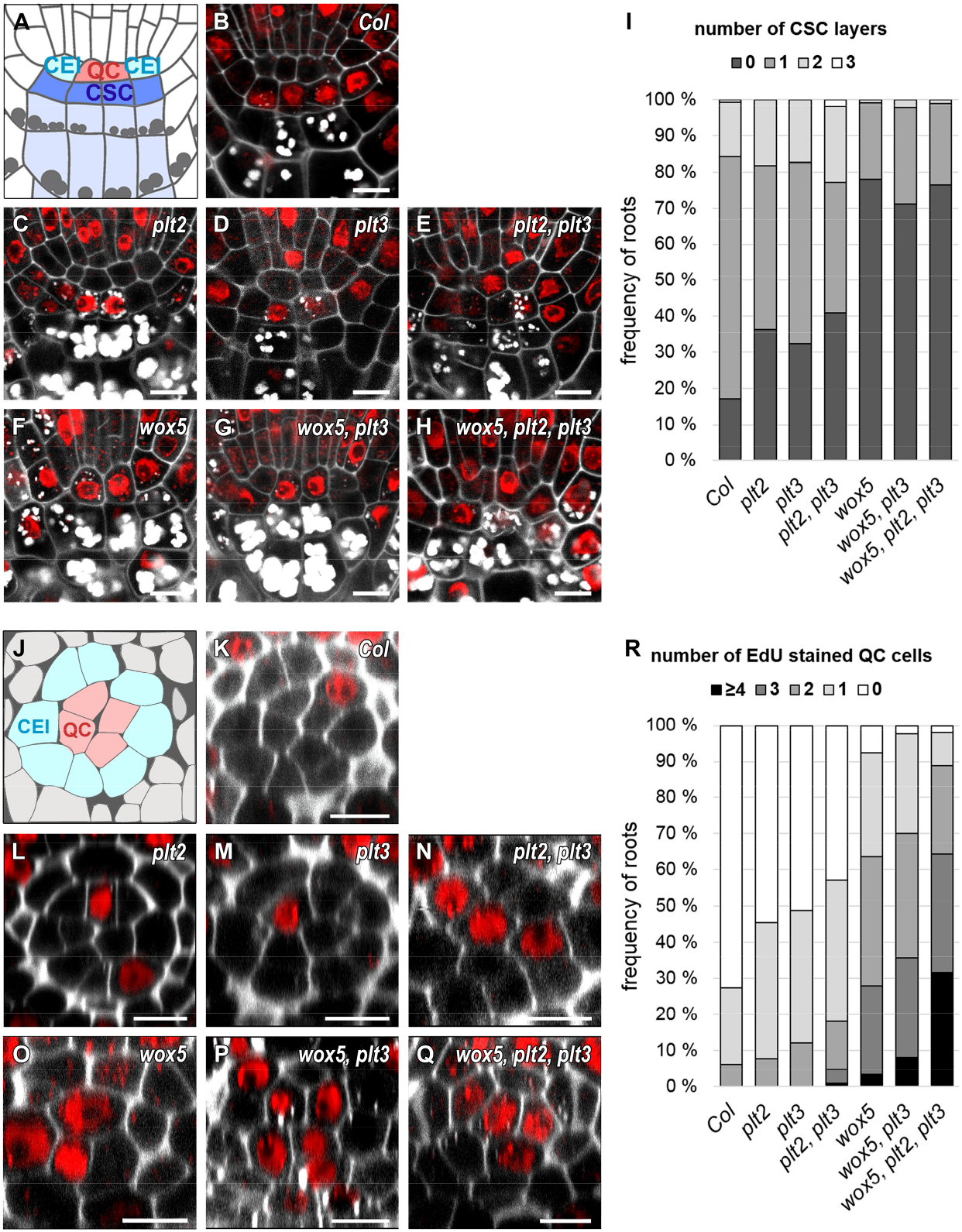
*plt* and *wox5* mutants show more CSC differentiation and QC divisions. **A**, Schematic representation of a longitudinal section of an *Arabidopsis* RM. QC cells are marked in red, CSCs are marked in dark blue, CCs in light blue. Combined mPSPI (grey) and EdU (red) staining for 24 hours (SCN staining) to analyse the CSC (**A**-**I**) and QC division phenotype (**J**-**R**) within the same roots are shown. **B**-**H**, Representative images of the SCN staining in *Col*, and the indicated single, double, and triple mutant roots. **I**, Analyses of the SCN staining for CSC phenotypes. Frequencies of roots showing 0, 1, 2, or 3 CSC layers are plotted as bar graphs. **J**, Schematic representation of a transversal section of an *Arabidopsis* RM. QC cells are marked in red, CEI initials are marked in turquoise. **R**, Analyses of the SCN staining for QC division phenotypes. Frequencies of roots showing 0, 1, 2, 3 or ≥4 dividing QC cells are plotted as bar graphs. Number of roots n = 77-146 from 2-5 independent experiments. QC = quiescent center, CSC = columella stem cell, CEI = cortex endodermis initial, SCN = stem cell niche, mPSPI = modified pseudo-Schiff propidium iodide, EdU = 5-ethynyl-2’-deoxyuridine, scale bars represent 5 μm.

WOX5 is necessary for CSC maintenance as loss of WOX5 causes their differentiation (Sarkar et al, 2007). In agreement with this, we found that the *wox5* mutants lack a starch-free cell layer in 78 % of analysed roots, indicating differentiation of the CSCs, compared to 17% in *Col* (Fig. 3A, B, F, I, Suppl. Table 7). In the *plt2* and *plt3* single mutants, the frequency of roots lacking a CSC layer increases to above 30 % (36 % and 32 %, respectively), and in the *plt2*, *plt3* double mutant to 41% (see Fig. 3C, D, E, I, Suppl. Table 7). Interestingly, the *wox5*, *plt3* double mutant as well as the *wox5*, *plt2*, *plt3* triple mutant show a frequency of differentiated CSCs comparable to the *wox5* single mutant (71 % and 77 %, respectively) (Fig. 3G, H, I, Suppl. Table 7). This data suggests that PLTs and WOX5 may act together in the same pathway to maintain CSC homeostasis, as there is no additive effect observable in the multiple mutant roots.

To analyse QC division phenotypes in detail, we quantified the number of EdU-stained cells in QC position in transversal optical sections. QC cells were identified by their position within the root SCN, as they are located directly distal to the stele initials and surrounded by the CEIs in a circular arrangement (Fig. 1A, J). In *Col*, 27 % of the analysed roots show at least one cell division in the QC within 24 hours (Fig. 3J, K, R, Suppl. Table 7), which is consistent with already published frequencies (Cruz-Ramírez et al, 2013). This frequency almost doubles to 45-50 % in the *plt2* and *plt3* single mutants and is even higher in the *plt2*, *plt3* double mutant (57 %) (Fig. 3L-N, R, Suppl. Table 7). Additionally, the *plt*-double mutant roots often show disordered QC regions with a disruption of the circular arrangement of cells surrounding the QC (Fig. 3N) which could be a result of uncontrolled divisions. *wox5* mutants show a disordered SCN accompanied by a high overall QC cell division frequency of at least one dividing QC cell in 92 % of roots (Fig. 3O, R) and on average more dividing QC cells per root (Suppl. Table 7). The number of dividing QC cells per root increases further in the *wox5*, *plt3* double mutant and is even higher in the *wox5*, *plt2*, *plt3* triple mutant; here, in one third of the roots all QC cells undergo cell division (Fig. 3P-R, Suppl. Table 7). Taken together, this data suggests an additive effect of PLT2, PLT3 and WOX5 regarding the QC-division phenotype, in accordance with our hypothesis that WOX5 and PLTs act in parallel pathways to maintain the quiescence of the QC.

Additionally, we quantified roots showing at least one aberrant periclinal cell division in the QC in longitudinal optical sections (Suppl. Fig. 1). Whereas the occurrence of these aberrant periclinal divisions in *Col* wild type roots is very rare (3 %), it increases in the *plt*-single mutants to 21 % and in *wox5* and *wox5*, *plt3* mutants to around 40 %. We found the most severe phenotypes in the *plt2*, *plt3* double and *wox5*, *plt2*, *plt3* triple mutants with an occurrence of periclinal QC-cell divisions in 53 % of the observed roots, indicating a synergistic regulatory role of PLTs in periclinal QC cell divisions (Suppl. Fig. 2B, Suppl. Table 8).

To visualise correlations of QC division and CSC differentiation, we combined the acquired data in 2D-plots in which the frequencies of the two phenotypes are color-coded (Fig. 4). This visualisation reveals a regular pattern for *Col* wild type roots, which peaks at one CSC-layer and no QC-divisions (Fig. 4A). The pattern of the *plt* single mutants is more irregular with a shift to less CSC-layers (indicating more differentiation) and more EdU-stained QC cells (indicating more QC divisions) compared to the wild type *Col* roots (Fig. 4B, C). The *plt2*, *plt3* double mutants have an additional maximum at a position showing no CSC layer and one divided QC cell, resulting in two phenotypic populations, one at a wild type-like position, the other showing a strong mutant phenotype (Fig. 4D). The 2D-pattern for the *wox5* mutant shifts to less CSC-layers and more QC-divisions with a maximum at no CSC-layers and two QC-divisions (Fig. 4E). The QC phenotype is more severe in the *wox5*, *plt3* double mutant towards more cell divisions and is even stronger in the *wox5*, *plt2*, *plt3* triple mutant which peaks at zero CSC layers and three QC-divisions (Fig. 4F, G). In summary, our data acquired by applying the novel SCN staining demonstrates that higher CSC differentiation correlates with a higher division rate in the QC, possibly to replenish missing stem cells by increased QC divisions.

**Figure 4 -.**
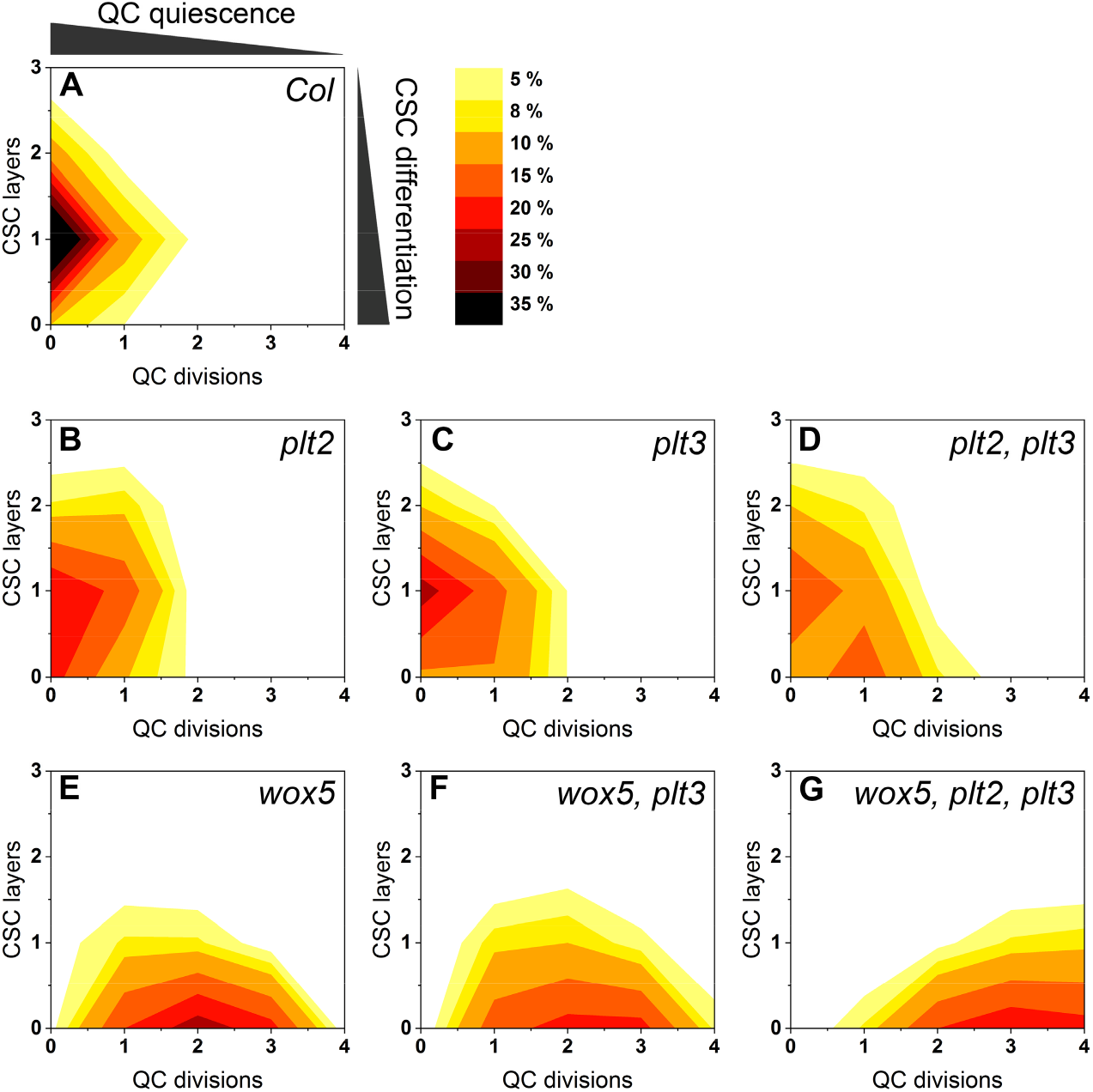
QC divisions correlate negatively with the number of CSC layers. The combined results of the SCN staining in Fig. 3 are shown as 2D plots to visualise the correlation of the CSC layer number and QC division. Number of CSC layers are shown on the y axis and the QC division phenotype is shown on the x axis. The darker the colour, the more roots show the respective phenotype (see colour gradient top right indicating the frequencies). *Col* wild type roots show one layer of CSCs and no EdU stained cells (no QC division) after 24 h EdU staining.

WOX5 and PLT3 are expressed and localise to overlapping domains in the SCN of the *Arabidopsis* root and based on our results regulate SCN maintenance together. To test for functionality of our generated reporter lines, we used the mVenus (mV) tagged WOX5 and PLT3 versions driven by their own endogenous promoters for rescue experiments in the respective mutant phenotypes in *Arabidopsis*. We observed a rescue of the *wox5* mutant expressing *pWOX5::WOX5-mV* and a rescue of the*plt3* mutant expressing *pPLT3::PLT3-mV* to almost wild type *Col* phenotypes, indicating that the labelling with mVenus did not or only very little influence WOX5 or PLT3 functionality (Suppl. Fig. 2, Suppl. Table 14). To our surprise, we observed PLT3 localisation in bright subnuclear structures, hereafter called NBs, in the PLT3-mV reporter line. Most frequently, we found PLT3 NBs in young, developing lateral root primordia (LRP) (Fig. 5A, Suppl. Movie 1) already at stages where PLT1 and PLT2 are not yet expressed (Du & Scheres, 2017). Importantly, we also observed PLT3 NBs in CSCs of some established primary roots, but never in QC cells (Fig. 5B-C’). To further examine the PLT3 NBs in a context, where no other PLTs are expressed, we used an estradiol-inducible system to control expression of FP-tagged PLT3 and WOX5 transiently in *Nicotiana benthamiana* leaf epidermal cells (Stahl et al, 2013). In agreement with our observations in *Arabidopsis*, we found that PLT3 mainly localises to NBs and to a lesser extend to the nucleoplasm (Fig. 6B). In co-expression experiments in *N. benthamiana*, we found that PLT3 recruits WOX5 to the same NBs, whereas on its own WOX5 remains homogenously localised within the nucleoplasm (Fig. 6G-G”, Suppl. Fig. 3A).

**Figure 5 -.**
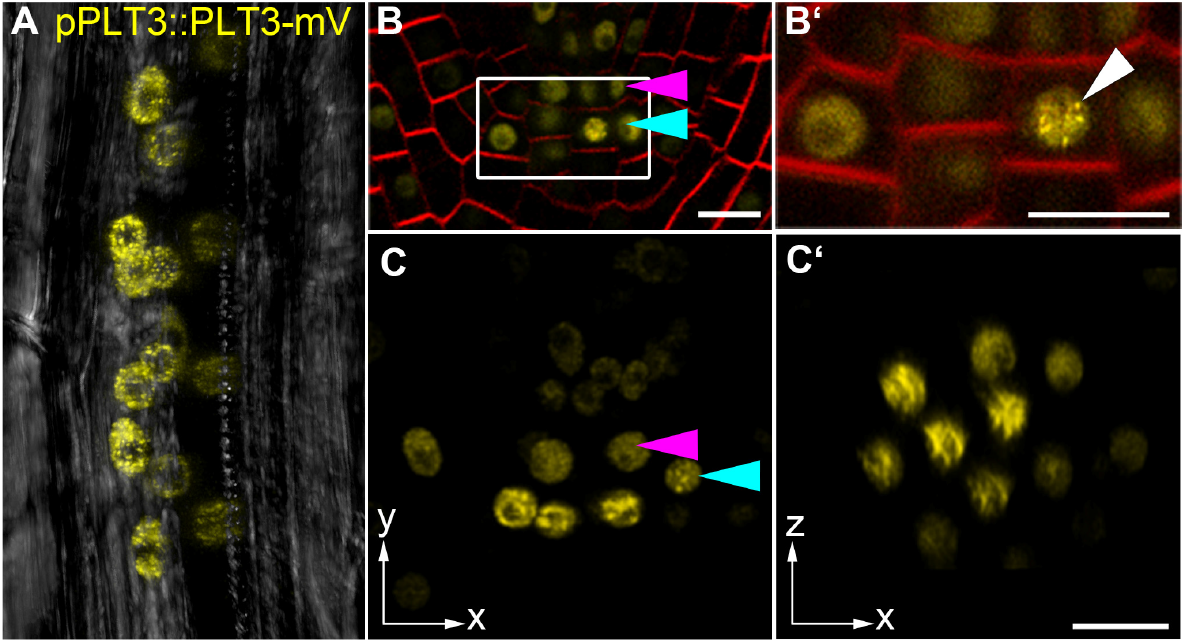
PLT3 localises to NBs in *Arabidopsis thaliana* LRPs and CSCs. A-C’, PLT3-mV expression driven by the PLT3 endogenous promoter in LRP (A) and primary root SCN (B-C’) in *plt3* mutant *Arabidopsis* roots. A, representative image of PLT3-mV expression (yellow) in an LRP showing the subnuclear localisation to NBs. Transmitted light image in grey. B, B’, SCN of an PLT3-mV expressing FM4-64-stained (red) *Arabidopsis* primary root. The magnification of the CSC layer (B’) shows the subnuclear localisation of PLT3 to NBs in a CSC. White arrowhead points at a NB. C, C’, SCN of an PLT3-mV expressing *Arabidopsis* primary root. NBs are visible in the CSC layer in C, also in the transversal view of the CSC layer (C’). Arrowheads in B and C point at the QC (magenta) and CSC (cyan) positions. mV = mVenus; LRP = lateral root primordium; SCN = stem cell niche; NBs = nuclear bodies; CSC = columella stem cell. Scale bars represent 10 μm.

**Figure 6 -.**
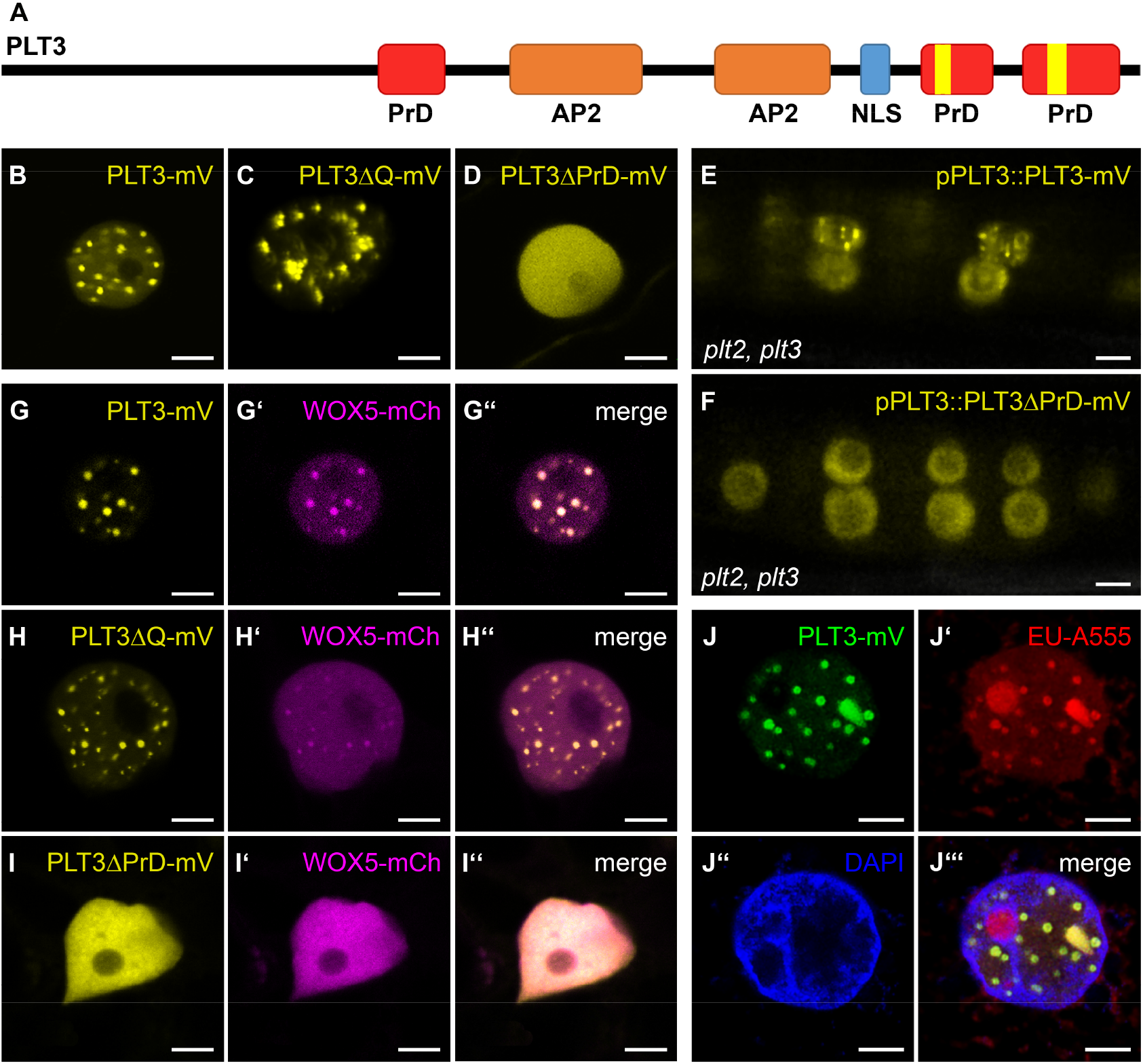
PLT3 PrD domains influence its subnuclear localisation. **A**, schematic representation of PLT3 protein domains. The areas in red are predicted prion-like domains (PrDs) and were deleted in PLT3ΔPrD-mV. The areas highlighted in yellow contain polyQ-stretches and were deleted in PLT3ΔQ-mV. Representative images of PLT3-mV (**B**), PLT3ΔQ-mV (**C**) and PLT3ΔPrD-mV (**D**) in transiently expressing *N. benthamiana* leaf epidermal cells. **E**, **F**, PLT3-mV (**E**) and PLT3ΔPrD-mV (**F**) expression driven by the PLT3 endogenous promoter in lateral root primordia of *plt2, plt3* double mutant *Arabidopsis* roots. **G-I’’**, Co-expression of PLT3-mV (**G**), PLTΔQ-mV (**H**) and PLT3ΔPrD-mV (**I**) with WOX5-mCh (**G’, H’, I’**) in transiently expressing *N. benthamiana* leaf epidermal cells. **J**-**J’’’**, Expression of PLT3-mV (**J**) in transiently expressing *N. benthamiana* leaf epidermal cells in combination with RNA staining with EU (18 h), visualised by click-reaction with Alexa Fluor^®^ 555 (**J’**) and a DNA staining with DAPI (**J’’**). mV = mVenus; PrD = prion-like domain; AP2 = APETALA2 domain; NLS = nuclear localisation signal; EU = 5-ethynyl-2’-uridine. Scale bars in (**B**-**J’’’**) represent 5 μm.

Next, we examined the protein domains putatively responsible for the localisation of PLT3 to NBs and found that the PLT3 amino acid (aa) sequence contains two glutamine (Q)-rich regions in the C-terminal part of the protein (see Fig. 6A). Proteins containing poly-Q stretches form aggregates or inclusions, a process often linked to pathological conditions in humans, such as Huntington’s disease (Scarafone et al, 2012). However, polyQ proteins can also convey diverse cellular functions like promotion of nuclear assemblies (e.g. the transcription initiation complex), formation of protein-protein complexes, recruitment of other polyQ-containing proteins (Atanesyan et al, 2012; Mikecz, 2009), and enhancement of the transcriptional activation potential of TFs (Schwechheimer et al, 1998; Gerber et al, 1994; Atanesyan et al, 2012). PolyQ domains were also found to be enriched in plant TFs (Kottenhagen et al, 2012).

Next, we tested if the polyQ-stretches in PLT3 are responsible for the subnuclear localisation and the recruitment of WOX5 to NBs. To this end, we deleted the polyQ domains of PLT3 and expressed the resulting PLT3ΔQ fused to mVenus transiently in *N. benthamiana*. We found that the subnuclear localisation and the recruitment of WOX5 did not change compared to the full-length PLT3 (see Fig. 6B, C, H-H’’). Therefore, we conclude that the polyQ domains in PLT3 are not, or at least not alone, responsible for the subnuclear localisation and translocation to NBs.

Apart from proteins with polyQ domains, many proteins that form concentration-dependent aggregates contain larger, intrinsically disordered regions (IDRs) with a low complexity similar to yeast prions (Cuevas-Velazquez & Dinneny, 2018). Prion-like proteins in *Arabidopsis* were first discovered by analysing protein sequences of 31 different organisms, identifying Q- and N-rich regions in the proteins to be sufficient to cause protein aggregation (Michelitsch & Weissman, 2000). Recently, the existence of more than 500 proteins with prion-like behaviour in *Arabidopsis* was reported (Chakrabortee et al, 2016) and the presence of prion-like domains (PrDs) in protein sequences is predictable with web-based tools (Lancaster et al, 2014). Therefore, we analysed the PLTs and WOX5 sequences using the PLAAC PrD prediction tool and found that PLT3 has three predicted PrDs in its aa sequence, two of them located at the C-terminus, containing the above described two polyQ-stretches (see Fig. 6A, Suppl. Fig. 3). PLT1, PLT2 and PLT4 also show predicted PrD domains, but PLT1 and PLT2 contain no and PLT4 shorter polyQ stretches within them. WOX5 does not show any predicted PrD domains, nor any polyQ stretches (Suppl. Fig. 3). To test the importance of the PrD domains in PLT3, we replaced the first PrD by a 27 aa linker (AAGAAGGAGGGAAAAAGGAGAAAAAGA) and deleted the C-terminally located PrDs. The resulting PLT3-version (PLT3ΔPrD) was fused to the mVenus FP and inducibly expressed in *N. benthamiana* epidermal cells. Here, we did not observe a localisation of PLT3ΔPrD-mVenus to NBs, but in contrast a homogenous distribution within the nucleus (Fig. 6D). In addition, upon co-expression of PLT3ΔPrD-mVenus with WOX5-mCherry, we observed that WOX5 was no longer recruited to NBs (Fig. 6I-I”). In line with this, we observed PLT3 NBs in developing *Arabidopsis* LRP expressing pPLT3::PLT3-mVenus, but no more NBs were found in a pPLT3::PLT3ΔPrD-mVenus expressing line (Fig. 6E, F). Based on these observations, we conclude that the PrD domains of PLT3 are responsible for its localisation to NBs and the recruitment of WOX5 to NBs. This is further supported by our observation that PLT3, in contrast to PLT1, 2 and 4, is found most frequently in NBs in transiently expressing in *N. benthamiana* correlating with the presence of PrD domains containing long polyQ-stretches in its aa sequence (Suppl. Fig. 3).

Proteins containing polyQ-stretches or PrDs are often involved in RNA binding, RNA processing and/or RNA compartmentalisation (Castilla-Llorente & Ramos, 2014; Alberti et al, 2009; Schomburg et al, 2001; Macknight et al, 1997; Sonmez et al, 2011; Fang et al, 2019). To test if PLT3 is involved in these processes, we performed an RNA-staining in *N. benthamiana* epidermal cells transiently expressing PLT3-mVenus with 5-ethynyl-2’-uridine (EU) (see Fig. 6J-J’’’). EU is incorporated into RNA during transcription, and we found that most of the stained RNA co-localises with the PLT3-mVenus NBs except for the EU-stained nucleolus (see Fig. 6J-J’’’). Based on these observations, we conclude that the PLT NBs act as important sites for the recruitment of RNA and other factors, including WOX5.

Because the WOX5 and PLT protein expression domains overlap in the SCN and PLT1, PLT2, PLT3 and PLT4 contain PrD domains, we asked whether PLTs and WOX5 can interact *in vivo*, especially considering the observed recruitment of WOX5 to PLT3 NBs. For this, we used fluorescence lifetime imaging microscopy (FLIM) to measure Förster resonance energy transfer (FRET) to analyse the potential protein-protein interaction of WOX5 and PLTs *in vivo*. To perform FLIM, we inducibly co-expressed WOX5-mVenus as donor together with individual PLTs-mCherry as acceptors for FRET in *N. benthamiana* leaf epidermal cells. The fluorescence lifetime of the donor fluorophore mVenus fused to WOX5 alone is 3.03 ± 0.07 ns. A reduction of fluorescence lifetime is due to Förster resonance energy transfer (FRET) of the two fluorophores in very close proximity (≤ 10 nm) caused by the direct interaction of the two observed proteins. When free mCherry is co-expressed as a negative control, the WOX5-mVenus mean fluorescence lifetime is not significantly decreased (3.00 ± 0.06 ns) (Fig. 7A, B, H, Suppl. Table 12). When WOX5-mVenus is co-expressed with PLT1-mCherry the fluorescence lifetime significantly decreases to 2.79 ± 0.11 ns, with PLT2-mCherry to 2.73 ± 0.12 ns, with PLT3-mCherry to 2.75 ± 0.17 ns and with PLT4-mCherry to 2.75 ± 0.24 ns, indicating FRET and hence direct protein-protein interactions (Fig. 7C-E, H, Suppl. Table 12). The observed interaction of WOX5 with PLT1-4 lead us to propose that they regulate SCN maintenance by the formation of complexes, either all together or in diverse compositions depending on the cell identity or their function. Interestingly, we observed a stronger lifetime decrease of WOX5-mVenus in the PLT3 NBs than in the nucleoplasm, indicating that the NBs function as main interaction sites of WOX5 with PLT3 (Fig. 7 I, J).

**Figure 7 -.**
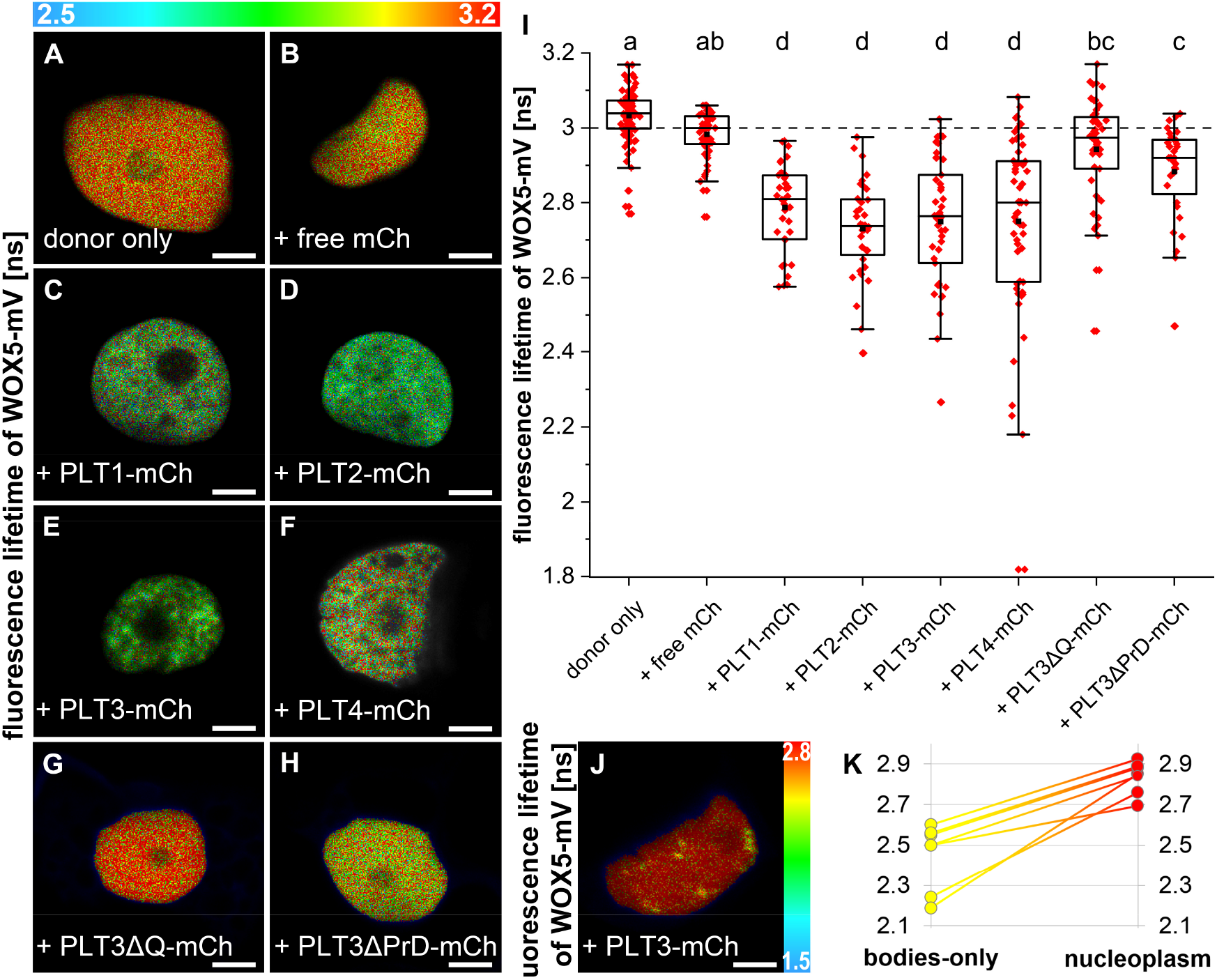
WOX5 can interact with PLTs. **A**-**H**, Fluorescence Lifetime Imaging (FLIM) results of transiently expressing *N. benthamiana* leaf epidermal cells. **A**-**H**, **J**, FLIM images of WOX5-mVenus (donor only) plus the indicated acceptors after a pixel-wise mono-exponential fit of the mVenus fluorescence signal. The fluorescence lifetime of WOX5-mVenus in ns is color-coded. Low lifetimes (blue) due to FRET indicate strong interaction of the two proteins and high lifetimes (red) indicate weaker or no interaction. Scale bars represent 5 μm. **I**, Fluorescence lifetimes in ns are summarized in combined jitter and box plots. The dashed line represents the fluorescence lifetime mean value of the WOX5-mV co-expressed with free mCh as negative control. After confirming normal distribution of the data by Kolmogorov-Smirnov testing, statistical analysis of samples was carried out by one-way ANOVA and Holm-Sidak post-hoc multiple comparisons test. Samples with identical letters do not show significant differences (α = 0.01; n ≥ 32). Box = 25-75 % of percentile, whisker = 1.5 interquartile range, − = median, ■ = mean value. **K**, seven individual nuclei showing nuclear bodies during co-expression of WOX5-mV and PLT3-mCh were analysed for WOX5-mV lifetime in the nuclear bodies or nucleoplasm separately. mCh = mCherry. mV = mVenus.

To address this, we measured the interaction between WOX5 and PLT3 in *Arabidopsis* roots via FLIM experiments in a translational line expressing WOX5-mVenus and PLT3-mCherry under control of their respective endogenous promoters. This resulted in the inevitable low protein concentration in comparison to the inducible system used in *N. benthamiana*. Probably due to this, we could not observe NBs in established root meristems of our *Arabidopsis* FLIM line and we could only measure a very small, albeit statistically significant, decrease in fluorescence lifetime from 2.97 ± 0.07 ns in the pWOX5::WOX5-mVenus (donor only FRET control) expressing roots to 2.94 ± 0.08 ns when pPLT3:PLT3-mCherry is co-expressed (Suppl. Fig. 4, Suppl. Table 13). In *Arabidopsis* seedlings, we only sometimes observed PLT3 NBs in the CSC layer of the root tip, but more frequently in young, developing LRP (Fig. 5), whereas in *N. benthamiana* we observed NBs in almost all cells. Therefore, we argue that the formation of the NBs is flexible because it is concentration dependent. In a transient *N. benthamiana* experiment, we could observe a correlation between the fluorescence intensity of nuclear PLT3-mVenus and the size and number of the NBs (Suppl. Fig. 5). NBs start to form after a certain intensity-threshold (Suppl. Fig. 5F) and their number decreases while their volume increases with overall rising fluorescence intensity (Suppl. Fig. 5A-G). A similar concentration-dependency has been previously described for proteins that undergo phase separation, in particular for liquid-liquid phase separation (LLPS) (McSwiggen et al, 2019). This mechanism separates membrane-free microdomains from the surrounding liquid and could potentially represent the underlying NB-forming process of PLTs. This process is possibly PrD-dependent as we observed less NB formation in the PrD-deletion variant of PLT3 (Fig.6 D, F).

Moreover, we asked if the PrD and poly-Q domains in PLT3 are required for protein-protein interaction with WOX5. To test this, we performed FLIM experiments with mCherry-tagged full-length PLT3, PLT3ΔQ and PLT3ΔPrD as acceptors and WOX5-mVenus as donor in *N. benthamiana*. Here, we observed that co-expression of the PLT3ΔQ and PLT3ΔPrD deletion variants did not lead to a significantly reduced fluorescence lifetime, and therefore no protein-protein interaction takes place in comparison to the full-length version (see Fig. 7E-H). This implies that PrD domains, containing the polyQ domains in PLT3, are necessary for the NB localisation, but also, notably, for protein complex-formation with WOX5.

Next, we asked if the PrD domains of PLT3 are also required for root SCN homeostasis. Therefore, we tested if the *plt2*, *plt3* double mutant phenotypes can be rescued by expressing full-length PLT3 or PLT3ΔPrD under control of the endogenous PLT3 promoter (Suppl. Fig. 2, Suppl. Table 14). We observed that full length PLT3 expression can rescue the *plt2*, *plt3* double mutant phenotype of more QC divisions and less CSC layers to the expected levels of *plt2* or *plt3* single mutant phenotypes (Suppl. Fig. 1D-F, H, J, K Suppl. Table 14). Importantly, PLT3ΔPrD could not rescue the *plt2*, *plt3* double mutant phenotype, suggesting that the PrD domains in PLT3 are indeed functionally relevant for SCN regulation and maintenance (Suppl. Fig. 2I-K, Suppl. Table 14).

In summary, our findings show that QC quiescence and CSC maintenance are mediated by mutual transcriptional regulation of PLTs and WOX5 as well as their direct protein-protein interaction and subnuclear partitioning to NBs due to PrDs.

## Discussion

In summary, we found that in agreement with a previous report (Sarkar et al, 2007), high *PLT* expression in the QC-region is promoted by WOX5, which again confines WOX5 to a defined and restricted number of cells within the QC region. In line with this, loss of PLTs leads to an expanded expression domain of *WOX5* and a reduced QC quiescence reflected in more QC divisions. Therefore, we could confirm that the control of QC quiescence and CSC maintenance is mediated by mutual transcriptional regulation of PLTs and WOX5 possibly by negative feedback loop regulation. As *WOX5* expression is normally limited to the QC, the question arises if, in absence of PLTs, either the *WOX5* expression domain expands to regions surrounding the QC or the QC region itself expands and therefore also the expression domain of *WOX5*. Interestingly, a previous report demonstrated that the expression of several QC markers is missing or highly reduced in *plt* mutants, suggesting that they fail to maintain an intact QC (Aida et al, 2004). Therefore, the expansion of the *WOX5* expression domain in the *plt* mutants is likely uncorrelated to the QC identity of these cells.

The observed higher frequency of cell divisions in the QC region of *wox5* mutants could be explained by the reduced expression of *PLTs*, which consequently negatively impacts QC quiescence but also by a PLT-independent pathway where WOX5 itself may have a positive effect on QC quiescence. Previous findings also suggest that WOX5 maintains QC quiescence through the repression of CYCD activity (Forzani et al, 2014). Considering our observation that PLT2, PLT3 and WOX5 show additive effects regarding the QC division phenotype, we propose a model in which WOX5 and PLTs could act in parallel pathways to maintain QC quiescence. The noted correlation of reduced QC quiescence and higher CSC differentiation that we could now show for individual roots by our newly introduced SCN staining could be necessary to replenish missing stem cells by QC divisions. This possible explanation is in agreement with the proposed function of the QC as a long-term stem cell reservoir, especially in case of stress or damage (Vilarrasa-Blasi et al, 2014).

For CSC homeostasis, PLTs and WOX5 may act together in the same pathway, possibly by complex formation, as there is no observable additive effect in the multiple mutant roots which is in agreement with previous findings (Ding & Friml, 2010). We show that WOX5 can directly interact with PLT1, PLT2, PLT3 and PLT4, which indicates that they regulate CSC maintenance by the formation of complexes, either all together or in diverse homo- or heteromeric compositions depending on cell identity or function.

In transient *N. benthamiana* experiments, PLT3 forms NBs and recruits WOX5 into them. The stronger drop of fluorescence lifetime in NBs compared to the nucleoplasm measured by FLIM implies that the PLT3 NBs function as major sites of protein-protein interaction with WOX5, which could be due to a favoured complex-formation within the NBs or due to transport of preformed WOX5/PLT3 complexes from the cytosol to NBs. We could observe PLT3 NBs in cells of the CSC layer of some *Arabidopsis* primary root tips, but never in the QC region. Nevertheless, PLT3 NBs were found more frequently in several cells of developing LRPs. LRPs are in a younger and less-determined stage than the primary root and the observed subnuclear localisation to NBs could represent a marker for the occurring determination and future cell differentiation. This agrees with the observed localisation of PLT3 to NBs in the CSCs in some of the primary roots. Here, PLT3 NBs could represent compartments for the recruitment of and interaction with WOX5 and possibly other factors involved in CSC fate determination and maintenance.

Furthermore, we found that the aa sequence of PLT3 comprises polyQ containing PrDs that are necessary for the localisation to NBs, complex formation with WOX5, and for SCN maintenance in the primary root meristem. Proteins containing polyQ-stretches or PrDs are often involved in RNA binding, RNA processing and/or RNA compartmentalisation (Castilla-Llorente & Ramos, 2014; Alberti et al, 2009; Schomburg et al, 2001; Macknight et al, 1997; Sonmez et al, 2011; Fang et al, 2019) and we could show that the PLT3 NBs indeed co-localise with RNA. Just as PLT3, FLOWERING CONTROL LOCUS A (FCA) is a PrD-containing protein (Chakrabortee et al, 2016) that localises to subnuclear structures (Fang et al, 2019). The FCA bodies separate from the cytosol by LLPS to provide compartments for RNA 3’-end processing factors (Fang et al, 2019). Similarly, PLT3 NBs could represent compartments for the recruitment of interacting factors and RNA for further processing, sequestration, or transportation. As PLT3 is a TF, the co-localising RNA could also represent newly transcribed RNA at the transcription sites where PLT3 binds to DNA, e.g. to the WOX5 promoter region (Shimotohno et al, 2018). The possible liquid-like nature of the PLT3 NBs will be an interesting subject for further studies investigating its putative phase separation properties. In this study, we showed that the PLT3 NB formation is concentration dependent, which is indicative for LLPS. This concentration-dependency of PLT3 NB formation could also explain the rare occurrence of PLT3 NBs at endogenous protein levels in the CSCs of the *Arabidopsis* RAM. Here, NB formation is possibly triggered only above a certain protein concentration threshold serving as a read-out for future cell fate. In the established distal root meristem, this is not continuously needed and therefore the protein concentration is mainly maintained below this threshold, so that often no NBs are formed. Consequently, the observed PLT3-FP fluorescence intensity in the CSCs can vary and is lower or higher at a given time (Fig. 1e and 5b and c).

Therefore, we propose that the regulation of QC quiescence and CSC maintenance are not only mediated by the mutual transcriptional regulation of PLT and WOX5, but also, importantly, by building protein complexes that are differentially localised to distinct nuclei within the SCN (see Fig. 8). The observed subnuclear localisation of PLT3 to NBs could represent a marker for the determination to future cell differentiation in the CSC layer.

**Figure 8 -.**
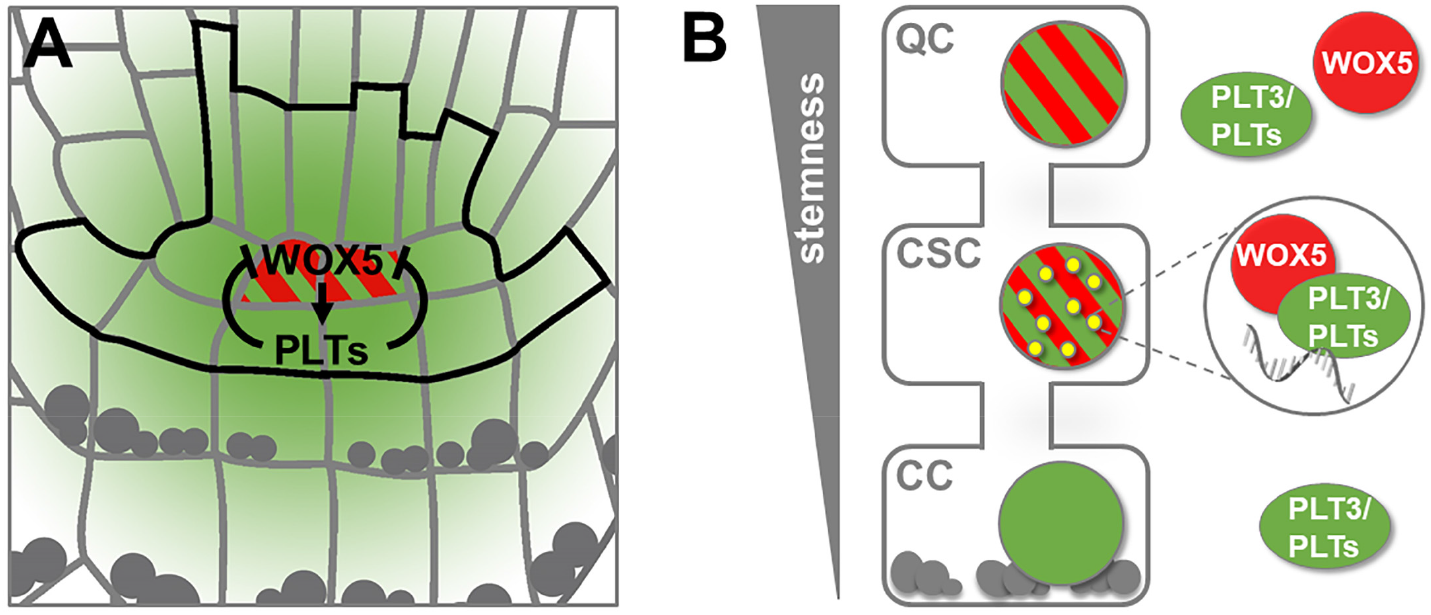
Model of PLT and WOX5 transcriptional regulation, interaction and subnuclear localisation during distal root stem cell maintenance. **A**, Transcriptional regulation of *WOX5* (red) and *PLT* (green) expression by negative feedback regulation in the *Arabidopsis* RAM. *WOX5* is expressed in the QC and promotes *PLT* expression, whereas *PLT* expression is restricting the *WOX5* expression domain to the QC position. **B**, Both WOX5 (red) and PLT3 (green) are present homogenously within the nuclei of the QC cells. WOX5 can move to the CSCs and is recruited there by PLT3 to NBs (yellow), where interaction takes place and RNA is present (grey in magnification). This maintains the stem cell character of the CSCs but already leads to a determination to subsequent CC fate. Gray dots represent starch granules.

Furthermore, the PrD and polyQ domains in PLT3 may act as an initial starting point to compartmentalise and partition WOX5 that has moved from the QC towards the CSC layer into NBs, possibly by a concentration dependent LLPS process. The observed sites could represent transcriptionally active sites for the regulation of target genes involved in CSC fate determination.

The dynamic compartmentalization to subcellular or subnuclear microdomains of proteins with intrinsically disordered, PrD and/or polyQ domains was shown to have severe effects, e.g., in human pathological disorders like Huntington’s disease (Scarafone et al, 2012). But also beneficial functions of prions that are responsible for some neurodegenerative diseases in mammals (Aguzzi et al, 2013; Kim et al, 2013) as a protein-based memory is highly discussed, as their self-replicating conformations could act as molecular memories to transmit heritable information (Bailey et al, 2004; Shorter & Lindquist, 2005). If this is also the case in plants remains to be determined. In general, the dynamic formation of subcellular structures could be necessary for a changing composition of assemblies in dependence of their functional status (Mikecz, 2009). The transition of these proteins between condensed and soluble forms requires high flexibility in their protein structure, which can be provided by flexible polyQ-stretches. Poly-Q domains are predominantly positioned at the surface of a protein, supporting the idea of their involvement in protein-protein interactions (Totzeck et al, 2017).

In *Arabidopsis*, PrD and poly-Q dependent compartmentalization could present a fast and reversible concentration-dependent regulatory mechanism (Cuevas-Velazquez & Dinneny, 2018), e.g. in case of PLT3 and WOX5 to determine CSC cell fate but probably also in other developmental contexts such as lateral root development. It remains to be determined if LLPS is the underlying mechanism of the observed subnuclear compartmentalisation of these key TFs involved in *Arabidopsis* root stem cell homeostasis and if also other processes necessary for determination of cell fates and stemness in *Arabidopsis* are regulated by this or similar dynamic processes.

## Methods

### Cloning

pWOX5::mVenus-NLS, pWOX5:: WOX5-mVenus, pPLT3::PLT3-mVenus, pPLT3::PLT3-mCherry, pPLT3::PLT3ΔPrD-mVenus and β-estradiol inducible PLT3ΔPrD-mVenus and β-estradiol inducible PLT3ΔPrD-mCherry were created by using the GreenGate cloning method (Lampropoulos et al, 2013). The internal *BsaI* restriction sites in the WOX5 promoter and WOX5 CDS were removed by PCR amplification of the sequences upstream and downstream of the *BsaI* sites with primer pairs whereof one primer has an altered nucleotide sequence at this site (Suppl. Table 1), followed by an overlap extension PCR to reconnect the gene fragments. The sequences upstream of the ATG start codon of WOX5 (4654 bp) and PLT3 (4494 bp) were used as promoter regions and were amplified by PCR and primers to add flanking *BsaI* restriction sites and matching overlaps for the GreenGate cloning system. Afterwards they were cloned into the GreenGate entry vector pGGA000 via *BsaI* restriction and ligation. The GreenGate promoter module carrying the β-estradiol inducible cassette was provided by (Denninger et al, 2019). The CDS of WOX5, PLT2, PLT3, PLT3ΔPrD and PLT4 as well as the FPs mVenus and mCherry were amplified by PCR using adequate primer pairs to add flanking *BsaI* restriction sites and matching overlaps for cloning into the GreenGate entry vectors pGGC000 (for CDS) and pGGD000 (for FPs) via *BsaI* restriction and ligation. All created entry vectors were confirmed by sequencing. The expression cassettes were created with a GreenGate reaction using pGGZ001 as destination vector. The correct assembly of the modules was controlled by sequencing. All module combinations used to construct the expression vectors can be found in Supplementary Table 3.

All other β-estradiol inducible constructs for *N. benthamiana* expression (free mCherry, WOX5-mVenus, WOX5-mCherry, PLT1-mVenus, PLT1-mCherry, PLT2-mCherry, PLT3-mVenus, PLT3-mCherry, PLT3ΔQ-mVenus, PLT3ΔQ-mCherry were created by Gateway cloning (Invitrogen^™^, Thermo Fisher Scientific Inc.). The CDS of WOX5, PLT1, PLT2, PLT3, PLT3ΔQ were amplified and cloned into pENTR/D-TOPO^®^. The Entry-vectors were confirmed by sequencing. The destination vector carrying the mVenus (pRD04) is based on pMDC7 (Curtis & Grossniklaus, 2003) which contains a β-estradiol inducible system for expression *in planta*. The mVenus was introduced via restriction/ligation C-terminally to the Gateway cloning site. The destination vector carrying the mCherry (pABindmCherry) was described before (Bleckmann et al, 2010). The expression vectors were created by LR-reaction of destination and entry vectors. Gateway expression vectors were verified by test digestion.

For the creation of the domain-deletion variants of PLT3 (PLT3ΔQ and PLT3ΔPrD), the CDS parts upstream and downstream of the desired sequence deletions were amplified with PCR and afterwards reconnected with overlap-PCR. The 27 aa linker (AAGAAGGAGGGAAAAAGGAGAAAAAGA) to replace the first PrD in PLT3ΔPrD was also introduced by overlap-PCR. All primer used for cloning can be found in Suppl. Table 1. For a list of the constructs produced in this study see Suppl. Table 3.

### Plant work

All *Arabidopsis* lines used in this study were in the Columbia (*Col-0*) background. The single mutants *wox5-1* and *plt3-1* have been described before(Galinha et al, 2007) (Suppl. table 4). The *plt2* (SALK_128164) and *wox5-1* (SALK038262) single mutants were provided by the *Arabidopsis* Biological Resource Center (ABRC, USA). The homozygous double and triple mutants were created by crossings (Supplementary table 4) and homozygous F3 genotypes were confirmed by PCR with appropriate primer pairs (Suppl. Table 2). The transgenic lines were created by floral dip as described before(Zhang et al, 2006) except for the published, transgenic *Col-0* lines with pPLT3::erCFP and pPLT3::PLT3-YFP (Galinha et al, 2007) constructs. They were crossed into the *wox5-1* mutant background. Homozygous lines were confirmed by genotyping and hygromycin selection. All plants for crossing, floral dips, genotyping and seed amplification were grown on soil in phytochambers under long day (16 h light/ 8 h dark) conditions at 21 °C. For microscopy *Arabidopsis* seeds were fume-sterilised (50 ml 13 % sodiumhypochlorite (v/v) +1 ml hydrochloric acid), imbedded in 0.2 % (w/v) agarose, stratified at 4 °C for 2 days and plated on GM agar plates (1/2 strength Murashige Skoog medium including Gamborg B5 vitamins, 1.2 % (w/v) plant agar, 1 % (w/v) sucrose, supplemented with 0.05 % (w/v) MES hydrate). *Arabidopsis* seedlings were grown for 5 days under continuous light at 21 °C and directly imaged afterwards.

### Cell wall and plasma membrane staining

For root imaging, the cell walls in *Arabidopsis* seedlings were stained by incubation in aqueous solutions of either 10 μM propidium iodide (PI) or 2.5 μM FM4-64 dye (Invitrogen^™^, Thermo Fisher Scientific Inc.). The staining solution was used as mounting medium for microscopy.

### *N. benthamiana* infiltration

For transient gene expression in *N. benthamiana*, the *Agrobacterium* strain GV3101::pMP50 was used as vector, carrying plasmids with the desired constructs and additionally either the helper plasmid p19 as silencing suppressor or the helper plasmid pSOUP that harbours a replicase needed for GreenGate vectors. Cultures were grown over night in 5 ml dYT-medium at 28 C on a shaker. The cultures were centrifuged for 10 min at 3345 g, the pellet was resuspended in infiltration medium (5 % (w/v) sucrose, 0.01 % (v/v) Silwet, 0.01 % (w/v) MgSO4, 0.01 % (w/v) glucose, 450 μM acetosyringone) to an optical density OD_600_ of 0.4 and cultures were incubated for one hour at 4 °C. The infiltration was done either with one single or with a combination of two different *Agrobacteria* cultures for co-expression of two constructs. A syringe without needle was used for the infiltration on the adaxial side of the leaves of well-watered *N. benthamiana* plants. For the expression of GreenGate constructs, an *Agrobacterium* strain carrying the p19 plasmid was co-infiltrated. The expression was induced 2-5 days after infiltration by applying an aqueous β-estradiol solution (20 μM β-Estradiol, 0.1 % (v/v) Tween^®^-20) to the adaxial leaf surface. Imaging or FLIM experiments were done 3 to 16 hours after induction, depending on the expression level.

### SCN staining

*Arabidopsis* seedlings were grown under continuous light for 5 days on GM agar plates without sucrose and then transferred on fresh plates containing additionally 7 μg/ml EdU to continue growing for 24 hours. Afterwards we performed an mPS-PI staining like described before(Truernit et al, 2008). Preliminary to the clearing step, the EdU-staining was performed. The permeabilisation of the cells and the subsequent staining of EdU-containing DNA with Alexa Fluor^®^ 488 was done as described in the Click-it^®^ EdU Imaging Kits from Invitrogen^™^ (Thermo Fisher Scientific Inc.) with adapted incubation times for *Arabidopsis* seedlings (permeabilisation for 1-2 h and click-reaction for 1 h). The click-reaction cocktail was prepared freshly with self-made solutions (Tris buffer with 50 mM Tris and 150 mM NaCl at pH 7.2-7.5; 4 mM CuSO4; 1.5 μM Alexa Fluor^®^ 488 picolyl azide;

50 mM ascorbic acid). The Alexa Fluor^®^ 488 picolyl azide (Thermo Fisher Scientific Inc.) was added from a 500 μM stock in DMSO. The ascorbic acid was added last from a freshly prepared aqueous 500 mM stock solution. After staining was done, the clearing step with chloralhydrate was performed like described beforeTruernit et al.

Images were acquired with a ZEISS LSM880 confocal microscope with imaging acquisition settings as stated below. z-stacks through the QC-region were recorded to obtain transversal views. To calculate the CSC phenotype, the number of CSC layers was counted in xy-images of each root. The QC-division phenotype is the number of EdU-Alexa Fluor^®^ 488-stained cells in the QC, which was counted in the cross-sectional images up to a maximal number of 4 stained QC cells. The phenotype frequencies of CSC differentiation and QC divisions (Fig. 3) where visualised in bar graphs with Excel (Microsoft Office 365 ProPlus, Microsoft Corporation). In order to correlate the two investigated phenotypes, we combined the CSC data and the QC-division data in 2D-plots. The combined QC/CSC-phenotype of every root was entered in a matrix with QC-divisions on the x- and CSC layers on the y-axis. 2D plots were created with Origin 2018b and 2020b (OriginLab Corporation).

### RNA staining

The RNA-staining in *N. benthamiana* epidermal cells was done on *N. benthamiana* leaves harboring a construct for a β-Estradiol inducible *PLT3-mVenus* expression. 5-ethynyl-2’-uridine (EU) was infiltrated in *N. benthamiana* leaves the day before staining. The expression of PLT3-mVenus was induced the next morning, 3 hours before fixation of the plant tissue. For fixation and permeabilisation of cells, pieces of the leaves were cut and treated with 4 % (w/v) paraformaldehyde and 0.5 % (v/v) TritonX-100 in PBS under vacuum for 1 h. The click-reaction of EU with Alexa Fluor^®^ 555 picolyl azide was performed similarly to the EdU-Alexa Fluor^®^ 488-staining described for the SCN staining in this article. A DAPI-counterstaining was carried out afterwards by incubating the leaf pieces in 0.1 μg/ml DAPI for 30 min. PBS was used as mounting medium for imaging.

### Microscopy

Imaging of *Arabidopsis thaliana* roots and *Nicotiana benthamiana* leaves was carried out with a ZEISS LSM780 or LSM880. Excitation and detection of fluorescent emission of fluorescent dyes was done as follows: DAPI was exited at 405 nm and emission was detected at 408-486 nm, Cerulean was excited at 458 nm and emission was detected at 460-510 nm; CFP was excited at 458 nm and emission was detected at 463-547 nm. mVenus was excited at 514 nm and emission was detected at 517-560 nm, or for co-expression with red dyes excited at 488 nm and detected at 500-560 nm. YFP was excited at 514 nm and emission was detected at 518-548 nm. Alexa Fluor^®^ 488 was excited at 488 nm and emission was detected at 490-560 nm. Alexa Fluor^®^ 555 was excited at 561 nm and emission was detected at 565-640 nm. PI was excited at 561 nm and emission was detected at 590-710 nm. FM4-64 was excited at 514 nm or 561 nm and emission was detected at 670-760 nm. mCherry was excited at 561 nm and emission was detected at 590-640 nm. Imaging of more than one fluorophore was done in sequential mode to avoid cross talk. The movie of pPLT3::PLT3-mVenus in a lateral root primordium was acquired with a MuViSPIM (Luxendo, Bruker) light sheet microscope equipped with a 40x/0.8 Nikon objective with a 1.5x tube lens on the detection axis to provide a 60x magnification.

### Image deconvolution

The microscope images in Fig. 5 a, c-c’ were deconvolved with Huygens 16.10.0p3 64b (Scientific Volume Imaging B.V.).

### Analyses of expression patterns and levels in *Arabidopsis*

For the comparison of relative fluorescence levels in the SCN of 5 day old *Arabidopsis* seedlings expressing either transcriptionally FP tagged PLT3 (pPLT3::erCFP) or translationally FP tagged *PLT3* (pPLT3::PLT3-YFP) driven by the endogenous PLT3 promoter in either the *Col* wild type or the *wox5-1* mutant, images of 9-16 roots per genotype were acquired with constant settings per FP. A ZEISS LSM880 confocal microscope was used. The mean fluorescence levels were measured with Fiji(Schindelin et al, 2012) in equally sized rectangular ROIs including the QC and CSC positions in the SCN. The thereby generated values were normalised to the *Col* mean fluorescence intensity and visualised in box and scatter plots created with Origin 2018b (OriginLab Corporation).

Images of the root tips of 5 day old *Arabidopsis* seedlings expressing *mVenus-NLS* driven by the endogenous WOX5 promoter in *Col* and *plt2* or *plt3-1* single mutants and the *plt2,plt3* double mutant were acquired. Additionally, z-stacks through the QC region of the roots were recorded to get a transversal view of the QC. The visualisation and counting of nuclei with *WOX5* expression (Fig. 2) was done with Imaris (version 9.1.2, Bitplane, Oxford Instruments plc). Box and scatter plots showing the number of expressing nuclei were created with Origin 2018b and 2020b (OriginLab Corporation). For the heat-map images, 10 acquired images were overlaid with Fiji(Schindelin et al, 2012) and the resulting fluorescence distribution was displayed with a 16-colors lookup table. To calculate the area of lateral *WOX5* expression in the QC region, a freehand-ROI surrounding the expressing cells was created in every image with Fiji (Schindelin et al, 2012). The ROI-areas were visualised in box and scatter plots, and all statistical analyses were carried by using Origin 2018b and 2020b (OriginLab Corporation).

### FLIM measurements

FLIM was performed either in *N. benthamiana* leaf epidermal cells expressing the desired gene combinations or in roots of 6-10 dag old *Arabidopsis* seedlings expressing *WOX5*-*mVenus* and *PLT3*-*mCherry* with their endogenous promoters. The FLIM measurements in *Arabidopsis* were performed in LRPs due to higher fluorescence levels and less movement during measurements compared to the RAM. mVenus-tagged proteins were always used as donor and mCherry-tagged proteins as acceptor for FRET. A ZEISS LSM 780 was used for the experiments equipped with a time correlated single-photon counting device (Hydra Harp 400, PicoQuant GmbH). The mVenus donor was excited with a linearly polarized diode laser (LDH-D-C-485) at 485 nm and a pulse frequency of 32 MHz. The excitation power was adjusted to 0.1-0.5 μW at the objective (C-Apochromat 40x/1.2 W Corr M27, ZEISS) for experiments in *N. benthamiana* and 1.5-2 μW for experiments in *Arabidopsis*. The higher laser power in *Arabidopsis* was needed due to lower fluorescence levels. τ-SPAD single photon counting modules with 2 channel detection units (PicoQuant GmbH) and a bandpass filter (534/30) were used to detect parallel and perpendicular polarized emission of the mVenus fluorescence. Images were acquired with a frame size of 256×256 pixel, a pixel dwell time of 12.6 μs and a zoom factor of 8. 40 to 60 frames were recorded in the *N. benthamiana* experiments, 80 frames in the experiments performed in *Arabidopsis*.

Fluorescent lifetimes were obtained by further analyses of the acquired data with SymPhoTime64 (PicoQuant GmbH). The instrument response function (IRF) of the microscope hardware is needed for fluorescence lifetime calculation to correct the systems-pecific internal time lag between laser pulse and data acquisition. The IRF was recorded preliminary to each experiment by time-correlated single photon counting (TCSPC) of an aqueous solution of erythrosine B in saturated potassium iodide. For data analysis of *N. benthamiana* experiments, an intensity threshold of 100-200 photons per pixel was applied to remove background fluorescence and a monoexponential fit was used. Due to low fluorescence intensities in *Arabidopsis* experiments, no threshold was applied to obtain the maximal possible photon number. In this case, a two-exponential fit was used to separate the mVenus fluorescence signal from the background fluorescence created by the plant tissue. This results in two lifetimes whereof one matches with the mVenus fluorescence lifetime of about 3 ns and the other representing the very short background lifetime of less than 0.4 ns. All data was obtained in at least two independent experiments. For visualisation of the lifetimes, box and scatter plots were created with Origin 2020b (OriginLab Corporation). Lifetime images of representative measurements were created with a pixel wise FLIM-fit in SymPhoTime64 (PicoQuant GmbH). The line graph showing the lifetime difference between the bodies and the nucleoplasm of WOX5-mVenus co-expressed with PLT3-mCherry was created using Excel (Microsoft Office 365 ProPlus, Microsoft Coorporation).

### Prediction of protein domains

The PrDs in the WOX5, PLT1, PLT2 and PLT3 aa sequences were predicted using the PLAAC application (Lancaster et al, 2014). The nuclear localisation signals (NLSs) of WOX5 and the studied PLT proteins were predicted using the cNLS Mapper(Kosugi et al, 2009) for WOX5 and PLT3 and SeqNLS (Lin & Hu, 2013) for PLT1 and PLT2.

### Figure assembly

All figures in this study were assembled using Adobe Photoshop (Adobe Inc.).

## Acknowledgements

We would like to acknowledge funding of R.C.B. by the Deutsche Forschungsgemeinschaft (DFG) through grant Sta1212/1-1 to Y.S. We thank Andrea Bleckmann for sharing the Greengate Cerulean construct, Renze Heidstra and Ben Scheres for sharing published *Arabidopsis* lines (PLT::PLT-YFP, PLT::CFP, PLT4 cDNA and *plt* mutants). We also thank the Center for Advanced Imaging (CAi) for technical assistance, and Rüdiger Simon and Peter Welters for critical discussion of the manuscript.

## Author contributions

R.C.B., V.I.S., A.A., A.D., L.C., and G.K.K. carried out experiments. Y.S. and R.C.B. designed the experiments, analysed and interpreted the data. S.W.P. contributed to FLIM data analyses. A.M. carried out light sheet imaging. Y.S. and R.C.B. wrote the manuscript. All authors commented on the manuscript.

## Material & correspondence

Correspondence and material requests should be addressed to Yvonne Stahl.

