## Supplementary material for "PLETHORA-WOX5 interaction and subnuclear localisation control *Arabidopsis* root stem cell maintenance": Suppl Figures

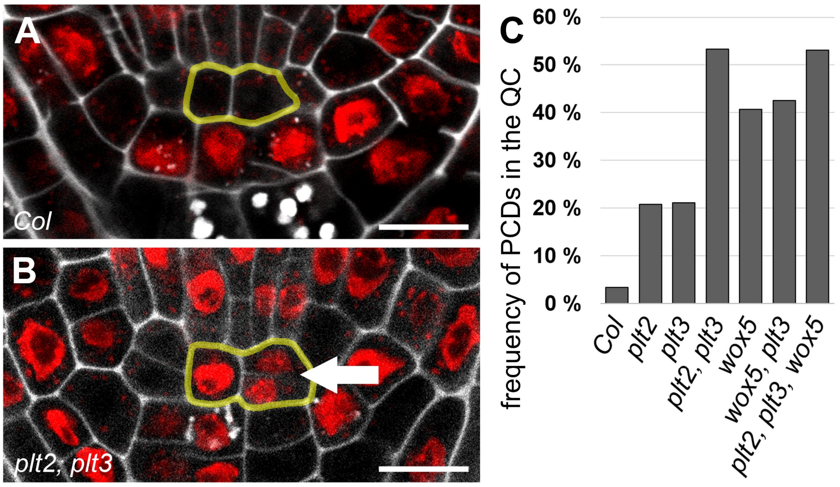


**Supplementary Figure 1 - *plt* and *wox5* mutants show more periclinal cell divisions in the QC.**

**A**, Representative figure of an *Arabidopsis* wildtype root SCN staining. **B,** Representative figure of an *Arabidopsis plt2, plt3* double mutant root SCN staining showing a periclinal cell division (PCD) in the QC (arrow). **A, B,** QC cells are outlined in yellow. Scale bars represent 10 µm. **C**, Analysis of the PCD phenotype. The frequency of roots (in percent) showing at least one PCD in the QC is plotted as a bar graph. Number of analysed roots n = 77-146. PCD = periclinal cell division.

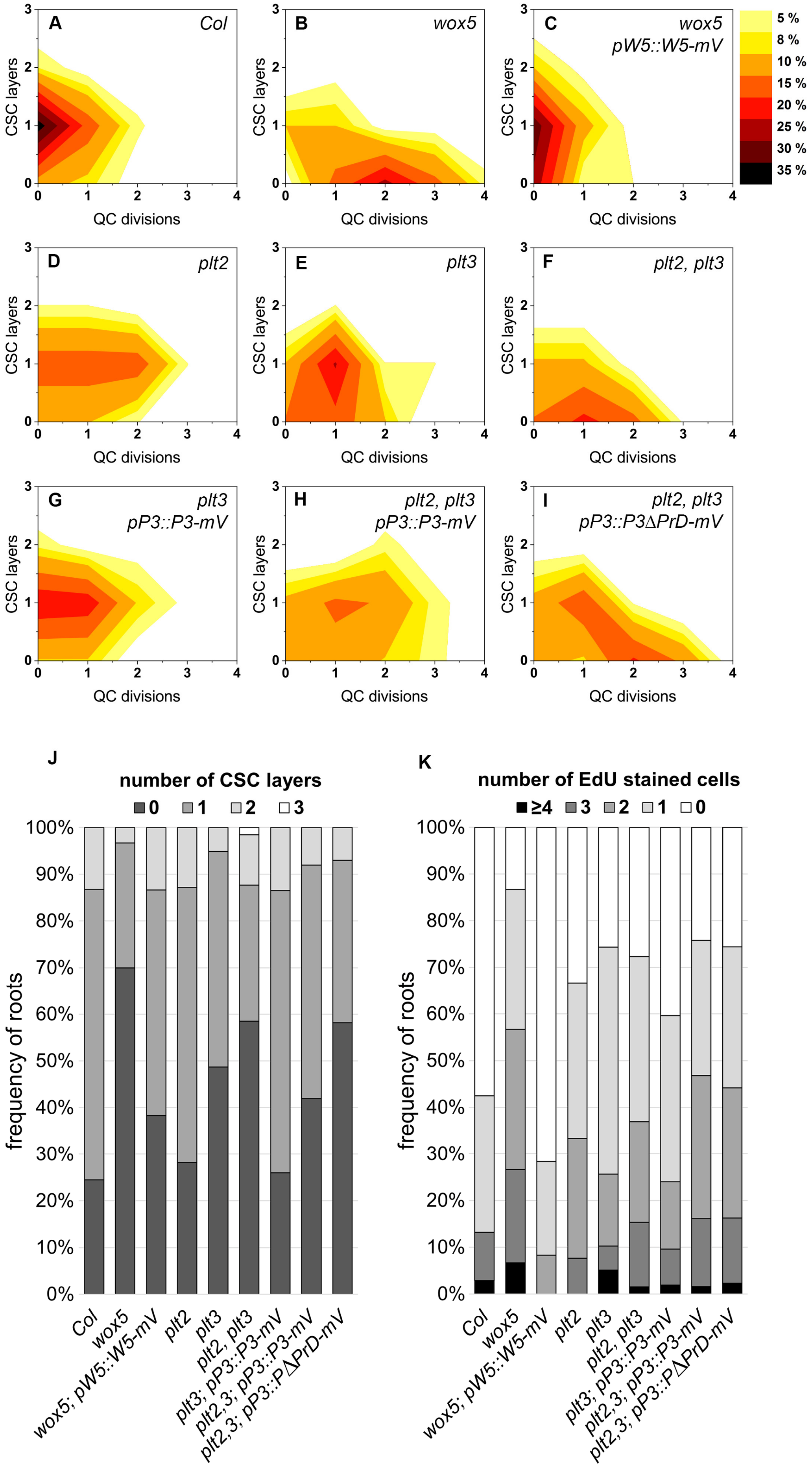
A
Col
B
wox5
C
wox5
pW5::W5-mV
5 %
8 %
10 %
15 %
20 %
25 %
30 %
35 %
D
plt2
E
plt3
F
plt2, plt3
G
plt3
pP3::P3-mV
H
plt2, plt3
pP3::P3-mV
I
plt2, plt3
pP3::P3ΔPrD-mV
CSC layers
QC divisions
J
number of CSC layers
0
1
2
3
frequency of roots
K
number of EdU stained cells
≥4
3
2
1
0
Col
wox5
wox5; pW5::W5-mV
plt2
plt3
plt2, plt3
plt3; pP3::P3-mV
plt2,3; pP3::P3-mV
plt2,3; pP3::PΔPrD-mV

10

**Supplementary Figure 2 - Mutant rescue experiments.**

SCN stainings were performed for two to five separate rescue experiments in *Arabidopsis thaliana* seedlings. The stainings were done in the indicated single and double mutant backgrounds expressing either WOX5-mV, PLT3-mV or PLT3ΔPrD-mV driven by their endogenous promoters as well as in *Col* wildtype. **A-I**, The combined results of the SCN staining are shown as 2D plots. Number of CSC layers are shown on the y axis and the QC division phenotype is shown on the x axis. The darker the colour, the more roots show the respective phenotype (see colour gradient on the right indicating the frequencies). **J, K**, Analyses of the SCN staining for CSC layer (**J**) or QC division (**K**) phenotypes. The frequencies of roots showing 0-3 CSC layers or 0-4 dividing QC cells are plotted as bar graphs. Number of analysed roots n = 30-106. EdU = 5-ethynyl-2′-deoxyuridine; CSC = columella stem cell; QC = quiescent centre; W5 = WOX5, P3 = PLT3.

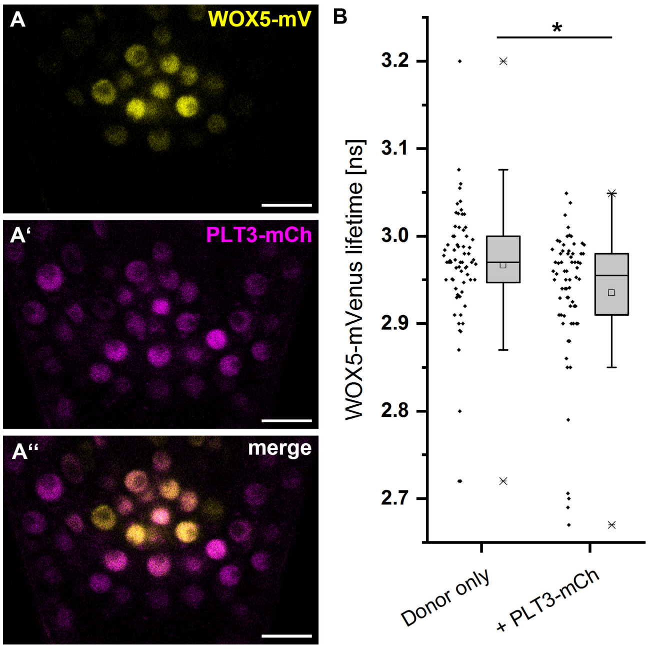


**Supplementary Figure 3 - PLT3-WOX5 interaction in the *Arabidopsis* root.**

**A-A'**, Representative image of the SCN in a lateral root of an *Arabidopsis* reporter line expressing WOX5-mV (**A**) and PLT3-mCh (**A'**) driven by their respective endogenous promoters. The TFs localize to overlapping domains (**A''**). Blue arrowheads mark QC cells, green arrowheads mark CSCs. Scalebars represent 10 µm. **B**, Fluorescence Lifetime Imaging (FLIM) results of experiments performed in *Arabidopsis thaliana* expressing either only WOX5-mV (donor-only) or both WOX5-mV and PLT3-mCh driven by their respective endogenous promoters. Donor fluorescence lifetimes in ns are summarized in combined scatter and box plots. The data is not normal distributed according to Kolmogorov-Smirnov test and therefore the non-parametric Krusak-Wallis ANOVA analysis with subsequent Dunns test was used to test for statistical significance ($\alpha = 0.01$). Asterisk marks statistically different samples. Number of measurements n = 67-68. mV = mVenus; mCh = mCherry; SCN = stem cell niche.

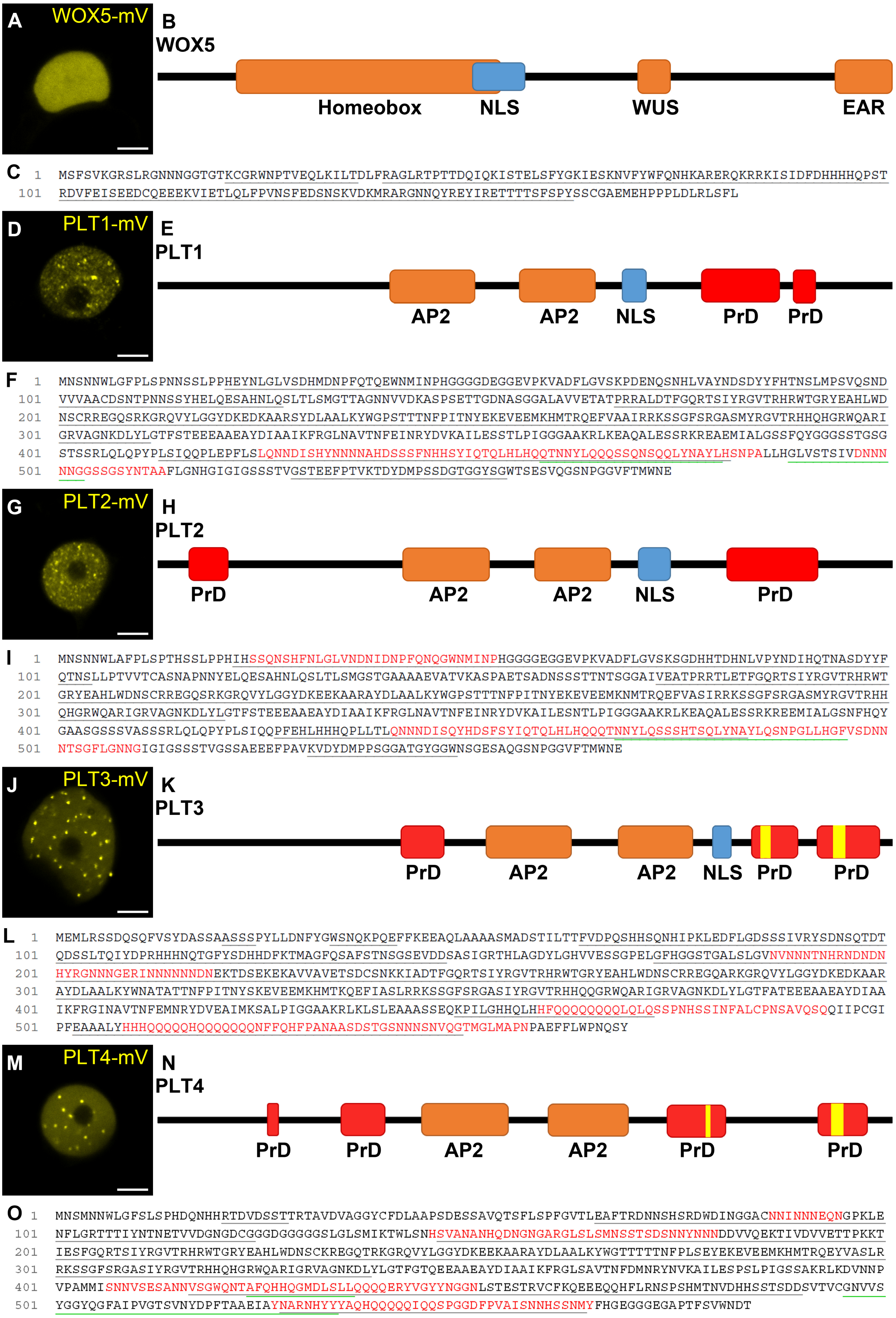
A
WOX5-mV
B
WOX5
Homeobox
NLS
WUS
EAR
C
1 MSFSVKGRSLRGNNNGGTGTKCGRWNPTVEQLKILTDLFRAGLRTPTTDQIQKISTELSFYGKIESKNVFYWFQNHKARERQKRRKISIDFDHHHHQPST
101 RDVFEISEEDCQEEEKVIETLQLFPVNSFEDSNSKVDKMRARGNNQYREYIRETTTTSFSPYSSCGAEMEHPPPLDLRLSFL
D
PLT1-mV
E
PLT1
AP2
AP2
NLS
PrD
PrD
F
1 MNSNNWLGFPLSPNNSSLPPHEYNLGLVSDHMDNPFQTQEWNMINPHGGGDEGGEVPKVADFLGVSKPDENQSNHLVAYNDSDYYFHTNSLMPSVQSND
101 VVVAACDSNTPNNSSYHELQESAHNLQSLTLSMGTTAGNNVVDKASPSETTGDNASGGALAVVETATPRRALDTFGQRTSIYRGVTRHRWTGRYEAHLWD
201 NSCRREGQSRKGRQVYLGGYDKEDKAARSYDLAALKYWGPSTTTNFPITNYEKEVEEMKHMTRQEFVAAIRRKSSGFSRGASMYRGVTRHHQHGRWQARI
301 GRVAGNKDLYLGTFSTEEEAAEAYDIAAIKFRGLNAVTNFEINRYDVKAILESSTLPIGGGAAKRLKEAQALESSRKREAEMIALGSSFQYGGGSSTGSG
401 STSSRLQLQPYPLSIQQPLEPFLSLQNNDISHYNNNNAHDSSSFNHHSYIQTQLHLHQQTNNYLQQSSQNSQQLYNAYLHSNPALLHGLVSTSIVDNNN
501 NNGGSSGSYNTAAFLGNHGIGIGSSSTVGSTEEFPTVKTDYDMPSSDGTGGYSGWTSESVQGSNPGGVFTMWNE
G
PLT2-mV
H
PLT2
PrD
AP2
AP2
NLS
PrD
I
1 MNSNNWLAFPLSPTHSSLPPHIHSSQNSHFNLGLVNDNIDNPFQNQGWNMINPHGGGGEGGEVPKVADFLGVSKSGDHHTDHNLVPYNDIHQTNASDYYF
101 QTNSLLPTVVTCASNAPNNYELQESAHNLQSLTLSMGSTGAAAAEVATVKASPAETSADNSSSTTNTSGGAIVEATPRRTLETFGQRTSIYRGVTRHRWT
201 GRYEAHLWDNSCRREGQSRKGRQVYLGGYDKEEKAARAYDLAALKYWGPSTTTNFPITNYEKEVEEMKNMTRQEFVASIRRKSSGFSRGASMYRGVTRHH
301 QHGRWQARIGRVAGNKDLYLGTFSTEEEAAEAYDIAAIKFRGLNAVTNFEINRYDVKAILESNTLPIGGGAAKRLKEAQALESSRKREEMIALGSNFHQY
401 GAASGSSSVASSSRLQLQPYPLSIQQPFEHLHHHQPLLTLQNNNDISQYHDSFSYIQTQLHLHQQQTNNYLQSSSHTSQLYNAYLQSNPGLLHGFVSDNN
501 NTSGFLGNNGIGIGSSSTVGSSAEEEFPAVKVDYDMPPSGGATGYGGWNSGESAQGSNPGGVFTMWNE
J
PLT3-mV
K
PLT3
PrD
AP2
AP2
NLS
PrD
PrD
L
1 MEMLRSSDQSQFVSYDASSAASSSPYLLDNFYGWSNQKPQEFFKEEAQLAAAASMADSTILTTFVDPQSHHSQNHIPKLEDFLGDSSSIVRYSDNSQTDT
101 QDSSLTQIYDPRHHHNQTGFYSDHHDFKTMAGFQSAFSTNSGSEVDDSASIGRTHLAGDYLGHVVESSGPELGFHGGSTGALSLGVNVNNNTNHRNDNDN
201 HYRGNNNGERINNNNNNDNEKTDSEKEKAVVAVETSDCSNKKIADTFGQRTSIYRGVTRHRWTGRYEAHLWDNSCRREGQARKGRQVYLGGYDKEDKAAR
301 AYDLAALKYWNATATTNFPITNYSKEVEEMKHMTKQEFIASLRRKSSGFSRGASIYRGVTRHHQQGRWQARIGRVAGNKDLYLGTFATEEEAAEAYDIAA
401 IKFRGINAVTNFEMNRYDVEAIMKSALPIGGAAKRLKLSLEAAASSEQKPILGHHQLHHFQQQQQQQQLQLQSSPNHSSINFALCPNSAVQSQQIIPCGI
501 PFEAAALYHHHQQQQQHQQQQQQQNFFQHFPANAASDSTGSNNNSNVQGTMGLMAPNPAEFFLWPNQSY
M
PLT4-mV
N
PLT4
PrD
PrD
AP2
AP2
PrD
PrD
O
1 MNSMNNWLGFSLSPHDQNHHRTDVDSSTTRTAVDVAGGYCFDLAAPSDESSAVQTSFLSPFGVTLEAFTRDNNSHSRDWDINGGACNNINNNEQNGPKLE
101 NFLGRTTTIYNTNETVVDGNGDCGGGDGGGGGSLGLSMIKTWLSNHSVANANHQDNGNGARGLSLSMNSSTSDSNNYNNNDDVVQEKTIVDVVETTPKKT
201 IESFGQRTSIYRGVTRHRWTGRYEAHLWDNSCKREGQTRKGRQVYLGGYDKEEKAARAYDLAALKYWGTTTTTNFPLSEYEKEVEEMKHMTRQEYVASLR
301 RKSSGFSRGASIYRGVTRHHQHGRWQARIGRVAGNKDLYLGTFGTQEEAAEAYDIAAIKFRGLSAVTNFDMNRYNVKAILESPSLPIGSSAKRLKDVNNP
401 VPAMMISNNVSESANNVSGWQNTAFQHHQGMDLSLLQQQQERYVGYYNGGNLSTESTRVCFKQEEEQQHFLRNSPSHMTNVDHHSSTSDDSVTVCGNVVS
501 YGGYQGFAIPVGTSVNYDPFTAAEIAYNARNHYYYAQHQQQQQIQQSPGGDFPVAISNNHSSNMYFHGEGGGEGAPTFSVWNDT

**Supplementary Figure 4 - Subnuclear localization and PrD prediction of WOX5, PLT1, PLT2, PLT3 and PLT4.**

**A, D, G, J, M**, (Sub-)nuclear localisation of WOX5-mV (**A**), PLT1-mV(**D**), PLT2-mV (**G**), PLT3-mV (**J**) and PLT4-mV (**m**) in transiently expressing *N. benthamina* epidermal cells. Scale bars represent 5 µm. **B, E, H, K, N**, schematic representation of WOX5 (**B**), PLT1 (**E**) PLT2 (**H**), PLT3 (**K**) and PLT4 (**N**) protein domains. The areas in red are predicted prion-like domains (PrDs), analysed using the PLAAC prediction tool. Yellow areas are polyQ stretches in the PLT3 and PLT4 amino acid sequence. **C, F, I, L, O**, Protein sequences of WOX5 (**C**) PLT1 (**F**), PLT2 (**I**), PLT3 (**L**) and PLT4 (**O**). The red highlighted sequences are the predicted prion-like domains (PrDs). mV = mVenus fluorescent protein; PrD = prion-like domain; EAR = Ethylene-responsive binding factor-associated repression domain; WUS = WUSCHEL box; AP2 = APETALA2 domain; NLS = nuclear localization signal.

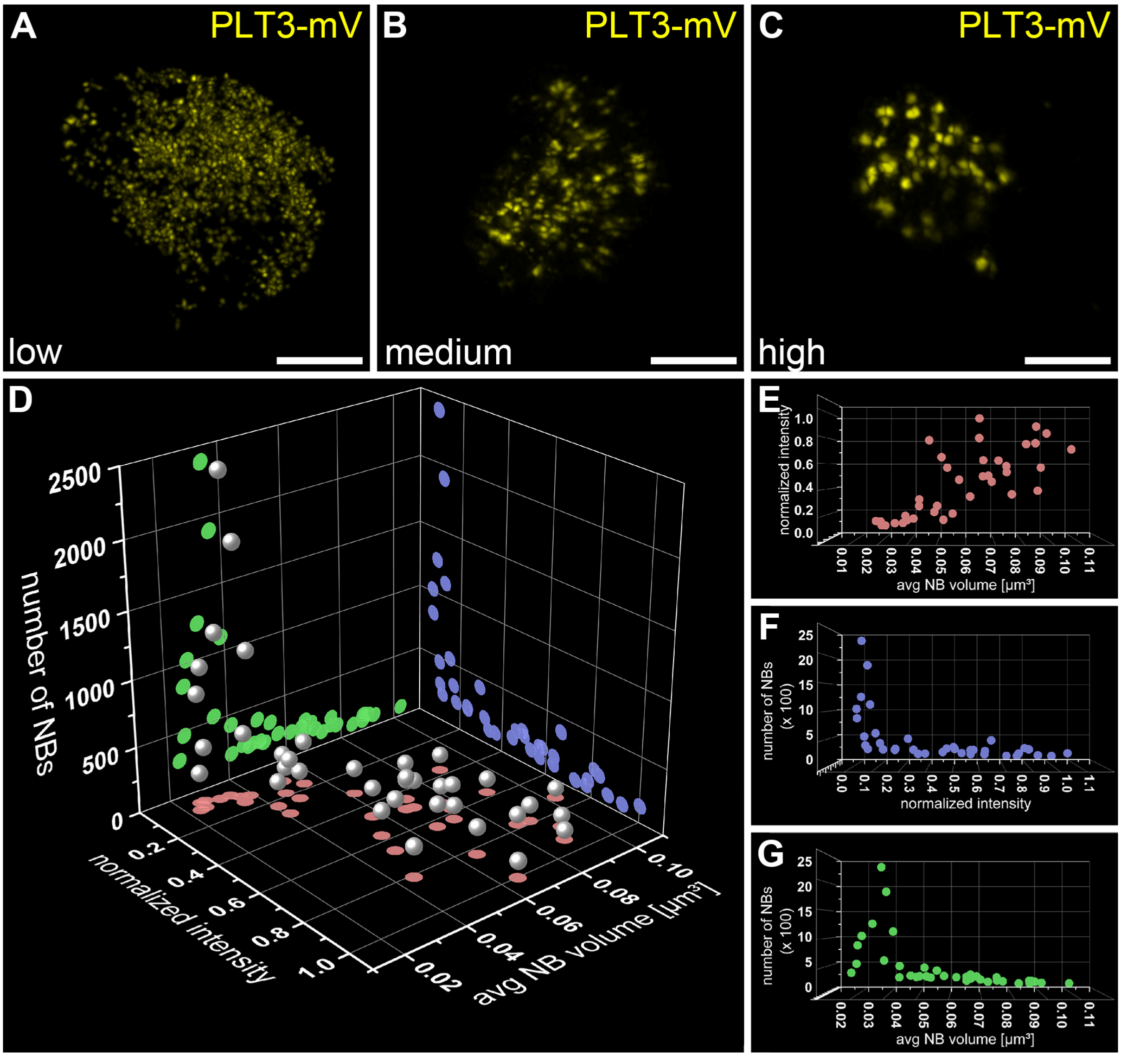


**Supplementary Figure 5 - Concentration dependency of PLT3 nuclear body formation.**

Representative images of low (**A**), medium (**B**), and high (**C**) PLT3-mVenus expressing nuclei in transiently expressing *N. benthamiana* leaf epidermal cells are shown. Scalebars represent 5 µm. **D-G,** analyses of intensities, numbers, and average volume of PLT3 NBs in individual nuclei, n = 37.

**Supplementary Movie 1 | Dynamic formation of nuclear bodies in a PLT3-mVenus expressing LRP.** The video shows a developing lateral root in an *Arabidopsis thaliana* plant expressing mVenus tagged PLT3 driven by the endogenous promoter (pPLT3::PLT3 mVenus) over 18 hours. Scale bar represents 25 µm.
