## Supplementary material for "PLETHORA-WOX5 interaction and subnuclear localisation control *Arabidopsis* root stem cell maintenance": Suppl Tables

#### Supplementary Table 1: List of cloning primers

| cloning system | gene ID | alias | primer name | orientation | Sequence 5´→ 3´ |
|---|---|---|---|---|---|
| GreenGate | | | **Promoter modules** | | |
| | AT3G11260 | pWOX5 | RD_GreenGate pWOX5 F | F | AAAGGTCTCAACCTAAAGACTTTTATCTACCAACTTCAAAAG |
| | | | RD_GreenGate pWOX5 R | R | AAAGGTCTCATGTTCGTTCAGATGTAAAG |
| | | | RD_GG pWOX5 BsaI a v2 F | F | AGAGACCAAATTATTTGGTTATATGGTAG |
| | | | RD_GG pWOX5 BsaI a v2 R | R | CTACCATATAACCAAATAATTTGGTCTCT |
| | | | RD_GreenGate pWOX5 BsaI b F | F | ATTACGATGTGAGAGCGCCTTCAACTTT |
| | | | RD_GreenGate pWOX5 BsaI b R | R | AAAGTTGAAGGCGCTCTCACATCGTAAT |
| | AT5G10510 | pPLT3 | RD_GG pPLT3 F V2 | F | AAAGGTCTCAACCTAATTTTAACGTATTCTTTC |
| | | | RD_GG pPLT3 R V2 | R | AAAGGTCTCATGTTAAACTTTCTTATAAAAACAATT |
| | | | **CDS modules** | | |
| | AT3G11260 | WOX5 | RD_GreenGate WOX5 F | F | AAAGGTCTCAGGCTTAATGTCTTTCTCCGTG |
| | | | RD_GreenGate WOX5 Mitte R | R | GACGTCGTGGTGGTTTCTCGAATATATT |
| | | | RD_GreenGate WOX5 Mitte F | F | AATATATTCGAGAAACCACCACGACGTC |
| | | | RD_GreenGate WOX5 R | R | AAAGGTCTCACTGAAAGAAAGCTTAATCG |
| | AT5G10510 | PLT3 | RD_GreenGate PLT3 F | F | AAAGGTCTCAGGCTTAATGGAGATGTTGAG |
| | | | RD_GreenGate PLT3 R | R | AAAGGTCTCACTGAGTAAGACTGATTAGGC |
| | | PLT3ΔPrD | RD_GreenGate PLT3 F | F | AAAGGTCTCAGGCTTAATGGAGATGTTGAG |
| | | | RD_PLT3ΔPrD1 CDS1 R | R | CCAGCTGCAACACCAAGTGACAAAG |
| | | | RD_PLT3ΔPrD1 linker F | F | CTTGGTGTTGCAGCTGGTGCTG |
| | | | RD_PLT3ΔPrD1 linker R | R | GTCTTCTCTGCTCCTGCGGCAG |
| | | | RD_PLT3ΔPrD1 CDS2 F | F | CGCAGGAGCAGAGAAGACAGATTCTG |
| | | | RD_GG PLT3ΔPrDs R | R | AAAGGTCTCACTGAGTGAAGTTGATGATGAC |
| | AT5G17430 | PLT4 | RD_GreenGate BBM F | F | AAAGGTCTCAGGCTTAATGAACTCGATGAAT |
| | | | RD_GreenGate BBM R | R | AAAGGTCTCACTGAAGTGTCGTTCCAAAC |
| | | | RD_GreenGate BBM Mitte F | F | ATTTACAATACCAACGAAACCGTTGTAGAT |
| | | | RD_GreenGate BBM Mitte R | R | ATCTACAACGGTTTCGTTGGTATTGTAAAT |
| Gateway | AT3G11260 | WOX5 | YS_WOX5 F CACC | F | CACCATGTCTTTCTCCGTGAAAGGTCGAAGCTTACG |
| | | | YS_WOX5 R -stop | R | AAGAAAGCTTAATCGAAGATCTAATGGC |
| | AT3G20840 | PLT1 | YS_PLT1 F CACC | F | CACCATGAATTCTAACAACTGGCTTGGCT |
| | | | YS_PLT1 r -stop | R | CTCATTCCACATAGTGAAAACACCACCAGGG |
| | AT1G51190 | PLT2 | YS_PLT2 F CACC | F | CACCATGAATTCTAACAACTGGCTCGCGTTCCCTCT |
| | | | YS_PLT2 R -stop | R | TTCATTCCACATCGTGAAAACACCTCCT |
| | AT5G10510 | PLT3 | YS_PLT3 F CACC | F | CACCATGGAGATGTTGAGGTCATCTGATCAGTCTCA |
| | | | YS_PLT3 R -stop | R | GTAAGACTGATTAGGCCAGAGGAAG |
| | | PLT3ΔQ | YS_PLT3 F CACC | F | CACCATGGAGATGTTGAGGTCATCTGATCAGTCTCA |
| | | | RD_PLT3 Seg1 R | R | GAGATGAGAAATGGTGAAGTTGATGATGAC |
| | | | RD_PLT3 Seg2 F | F | CTTCACCATTTCTCATCTCCTAATCACAGTAGC |
| | | | RD_PLT3 Seg2 R | R | GAAGAAGTTGTGGTGGTGGTAAAGAGCAG |
| | | | RD_PLT3 Seg3 F | F | CACCACCACAACTTCTTCCAGCATTTTCC |
| | | | YS_PLT3 R -stop | R | GTAAGACTGATTAGGCCAGAGGAAG |

#### Supplementary Table 2: List of genotyping primers

| gene ID | alias | primer name | orien-tation | Sequence 5´→ 3´ |
|---|---|---|---|---|
| AT3G11260 | *wox5-1* | GK_WOX5 F | F | AAACAGTTGAGGACTTTACATCTGA |
| | | WOX5 R | R | CGGATAATATGTCATAATTCAAAAT |
| AT5G10510 | *plt3-1* | GK_PLT3L | F | TTGTGATTTGCCATTGACTAAAGGT |
| | | GK_PLT3R | R | GAAAACAGTCCAATGGTCTCACATC |
| AT1G51190 | *plt2* | RD_plt2 SALK128164 neu L | F | GCTTAAATAGATATATGGTCATGCTTATATTC |
| | | RD_plt2 SALK128164 neu R | R | CAAGAAGACTCCAGCCGATC |
| | | SALK LBB1V2 | | AAACCAGCGTGGACCGCTTGCTGCAACTCT |

#### Supplementary Table 3: List of expression vectors created in this study

| cloning | constructs | promoter | N-tag | CDS | C-tag | term-inator | plant sel. marker | destination Vector | bacterial sel. | plasmid ID |
|---|---|---|---|---|---|---|---|---|---|---|
| | | module A | module B | module C | module D | module E | module F | module Z | | |
| GreenGate | **pWOX5:: mVenus-NLS** | WOX5 promoter | Ω-element (pGGB002) | mVenus | linker-NLS (pGGD007) | tUBQ10 (pGGE009) | BASTA (pGGF002) | pGGZ001 | Spec | pVS10 |
| | **pWOX5:: WOX5-mV** | WOX5 promoter | Ω-element (pGGB002) | WOX5 | mVenus | tUBQ10 (pGGE009) | Hyg (pGGF005) | pGGZ001 | Spec | pRD48 |
| | **pPLT3:: PLT3-mV** | PLT3 promoter | Ω-element (pGGB002) | PLT3 | mVenus | tUBQ10 (pGGE009) | Hyg (pGGF005) | pGGZ001 | Spec | pRD73 |
| | **pPLT3:: PLT3-mCh** | PLT3 promoter | Ω-element (pGGB002) | PLT3 | mCherry | tUBQ10 (pGGE009) | - | pGGM000 | Kan | pRD83 |
| | **pWOX5:: WOX5-mV** | WOX5 promoter | Ω-element (pGGB002) | WOX5 | mVenus | tUBQ10 (pGGE009) | Hyg (pGGF005) | pGGN000 | Kan | pRD84 |
| | **pPLT3::PLT3-mCh pWOX5::WOX5-mV** | pRD83 + pRD84 | | | | | | pGGZ001 | Spec | pRD89 |
| | **pPLT3:: PLT3ΔPrD-mV** | PLT3 promoter | Ω-element (pGGB002) | PLT3 ΔPrD | mVenus | tUBQ10 (pGGE009) | Hyg (pGGF005) | pGGZ001 | Spec | pRD125 |
| | **inducible PLT3ΔPrD-mV** | Ubi-XVE oLexA-35S | Ω-element (pGGB002) | PLT3 ΔPrD | mVenus | tUBQ10 (pGGE009) | Hyg (pGGF005) | pGGZ001 | Spec | pRD106 |
| | **inducible PLT3ΔPrD-mCh** | Ubi-XVE oLexA-35S | Ω-element (pGGB002) | PLT3 ΔPrD | mCherry | tUBQ10 (pGGE009) | Hyg (pGGF005) | pGGZ001 | Spec | pRD138 |
| | **inducible PLT2-mV** | Ubi-XVE oLexA-35S | Ω-element (pGGB002) | PLT2 | mVenus | tUBQ10 (pGGE009) | Hyg (pGGF005) | pGGZ001 | Spec | pRD102 |
| | **inducible PLT4-mV** | Ubi-XVE oLexA-35S | Ω-element (pGGB002) | PLT4 | mVenus | tUBQ10 (pGGE009) | Hyg (pGGF005) | pGGZ001 | Spec | pRD150 |
| | **inducible PLT4-mCh** | Ubi-XVE oLexA-35S | Ω-element (pGGB002) | PLT4 | mCherry | tUBQ10 (pGGE009) | Hyg (pGGF005) | pGGZ001 | Spec | pRD151 |
| Gateway | **inducible WOX5-mV** | Ubi-XVE oLexA-35S | - | WOX5 | mVenus | T3A | Hyg | pRD04 | Spec | pRD26 |
| | **inducible WOX5-mCh** | Ubi-XVE oLexA-35S | - | WOX5 | mCherry | T3A | Hyg | pABind mCherry | Spec | pFB02 |
| | **inducible PLT1-mV** | Ubi-XVE oLexA-35S | - | PLT1 | mVenus | T3A | Hyg | pRD04 | Spec | pRD56 |
| | **inducible PLT1-mCh** | Ubi-XVE oLexA-35S | - | PLT1 | mCherry | T3A | Hyg | pABind mCherry | Spec | pRD144 |
| | **inducible PLT2-mCh** | Ubi-XVE oLexA-35S | - | PLT2 | mCherry | T3A | Hyg | pABind mCherry | Spec | pRD147 |
| | **inducible PLT3-mV** | Ubi-XVE oLexA-35S | - | PLT3 | mVenus | T3A | Hyg | pRD04 | Spec | pRD25 |
| | **inducible PLT3-mCh** | Ubi-XVE oLexA-35S | - | PLT3 | mCherry | T3A | Hyg | pABind mCherry | Spec | pFB06 |
| | **inducible PLT3ΔQ-mV** | Ubi-XVE oLexA-35S | - | PLT3 ΔQ | mVenus | T3A | Hyg | pRD04 | Spec | pRD57 |
| | **inducible PLT3ΔQ-mCh** | Ubi-XVE oLexA-35S | - | PLT3 ΔQ | mCherry | T3A | Hyg | pABind mCherry | Spec | pRD81 |

**Supplementary Table 4: *Arabidopsis* mutants and transgenic lines**

| gene ID | alias | reference |
|---|---|---|
| AT3G11260 | *wox5-1* | SALK038262 (ABRC) |
| AT5G10510 | *plt3-1* | Galinha *et al*. (2007) |
| AT1G51190 | *plt2* | SALK_128164 (ABRC) |
| AT1G51190, AT5G10510 | *plt2, plt3* | this study, crossing of *plt2* and *plt3-1* |
| AT3G11260, AT5G10510 | *wox5, plt3* | this study, crossing of *wox5-1* and *plt3-1* |
| AT3G11260, AT1G51190, AT5G10510 | *wox5, plt2, plt3* | this study, crossing of *plt2, plt3* and *wox5, plt3* |
| AT5G10510 | pPLT3::erCFP (*Col-0*) | Galinha *et al*. (2007) |
| AT5G10510 | pPLT3::erCFP (*wox5-1*) | this study, crossing of pPLT3::erCFP (*Col-0*) and *wox5-1* |
| AT5G10510 | pPLT3::PLT3-YFP (*Col-0*) | Galinha *et al*. (2007) |
| AT5G10510 | pPLT3::PLT3-YFP (*wox5-1*) | this study, crossing of pPLT3::PLT3-YFP (*Col-0*) and *wox5-1* |
| AT5G10510 | pPLT3::PLT3-mVenus (*plt2, plt3*) | this study |
| AT5G10510 | pPLT3::PLT3ΔPrD-mVenus (*plt2, plt3*) | this study |
| AT3G11260 | pWOX5::mVenus-NLS (*Col-0*) | this study |
| AT3G11260 | pWOX5::mVenus-NLS (*plt2*) | this study |
| AT3G11260 | pWOX5::mVenus-NLS (*plt3-1*) | this study |
| AT3G11260 | pWOX5::mVenus-NLS (*plt2, plt3*) | this study |

**Supplementary Table 5: Intensity values of transcriptional and translational FP tagged PLT3 expression experiments in *Col-0* and *wox5* related to Figure 1.**

| root # | pPLT3::erCFP fluorescence intensity [%] | | pPLT3::PLT3-YFP fluorescence intensity [%] | |
|---|---|---|---|---|
| | ***Col-0*** | ***wox5*** | ***Col-0*** | ***wox5*** |
| **1** | 84 | 65 | 96 | 69 |
| **2** | 108 | 67 | 121 | 31 |
| **3** | 81 | 78 | 73 | 72 |
| **4** | 93 | 48 | 68 | 64 |
| **5** | 101 | 46 | 68 | 38 |
| **6** | 95 | 65 | 95 | 109 |
| **7** | 113 | 52 | 109 | 75 |
| **8** | 128 | 53 | 72 | 39 |
| **9** | 82 | 58 | 78 | 79 |
| **10** | 71 | | 127 | 30 |
| **11** | 105 | | 164 | 63 |
| **12** | 139 | | 84 | 39 |
| **13** | | | 145 | 66 |
| **14** | | | | 55 |
| **15** | | | | 56 |
| **16** | | | | 99 |
| **mean** | **100** | **59** | **100** | **62** |
| **st. dev.** | **20.10** | **10.39** | **31.13** | **23.01** |

**Supplementary Table 6: Quantification of transcriptional mVenus tagged WOX5 expression in the QC region of *Col* and *plt2, plt3* related to Figure 2.**

| root # | number of cells with WOX5 expression | | lateral area of WOX5 expression in the root tip [µm²] | |
|---|---|---|---|---|
| | ***Col*** | ***plt2, plt3*** | ***Col*** | ***plt2, plt3*** |
| **1** | 7 | 14 | 432.03 | 719.28 |
| **2** | 4 | 9 | 345.88 | 886.24 |
| **3** | 7 | 13 | 393.67 | 861.28 |
| **4** | 7 | 13 | 400.62 | 882.48 |
| **5** | 8 | 12 | 346.86 | 548.73 |
| **6** | 6 | 11 | 430.11 | 736.44 |
| **7** | 8 | 14 | 467.05 | 793.77 |
| **8** | 6 | 12 | 512.33 | 693.18 |
| **9** | 7 | 13 | 444.99 | 948.22 |
| **10** | 9 | 14 | 474.60 | 747.27 |
| **mean** | **6.90** | **12.50** | **424.81** | **781.69** |
| **st. dev.** | **1.30** | **1.50** | **51.26** | **111.59** |

**Supplementary Table 7: Average QC and CSC phenotypes related to Figure 3.**

| Genotype | average QC cell-divisions per root (± st. dev.) | average CSC layers per root (± st. dev.) | number of analyzed roots |
|---|---|---|---|
| ***Col*** | 0.34 ± 0.59 | 0.99 ± 0.59 | 146 |
| ***plt2*** | 0.53 ± 0.64 | 0.82 ± 0.72 | 77 |
| ***plt3*** | 0.61 ± 0.64 | 0.85 ± 0.69 | 121 |
| ***plt2, plt3*** | 0.81 ± 0.88 | 0.84 ± 0.82 | 105 |
| ***wox5*** | 1.87 ± 0.98 | 0.23 ± 0.44 | 118 |
| ***wox5, plt3*** | 2.11 ± 0.98 | 0.31 ± 0.51 | 87 |
| ***wox5, plt2, plt3*** | 2.83 ± 1.04 | 0.24 ± 0.46 | 98 |

**Supplementary Table 8: Percentage of periclinal cell divisions in the QC shown in Supplementary Figure 2.**

| Genotype | periclinal QC cell divisions [%] | number of analyzed roots |
|---|---|---|
| *Col* | 3 | 146 |
| *plt2* | 21 | 77 |
| *plt3* | 21 | 121 |
| *plt2, plt3* | 53 | 105 |
| *wox5* | 41 | 118 |
| *wox5, plt3* | 43 | 87 |
| *wox5, plt2, plt3* | 53 | 98 |

**Supplementary Table 9: Average QC and CSC phenotypes of rescue experiments**

| Genotype | average QC cell-divisions per root (± SD) | average CSC layers per root (± SD) | number of analyzed roots |
|---|---|---|---|
| ***Col*** | 0.58 ± 0.79 | 0.89 ± 0.60 | 106 |
| ***wox5*** | 1.77 ± 1.12 | 0.33 ± 0.54 | 30 |
| ***wox5 pWOX5::WOX5-mV*** | 0.43 ± 0.66 | 0.88 ± 0.65 | 51 |
| ***plt2*** | 1.08 ± 0.94 | 0.85 ± 0.62 | 39 |
| ***plt3*** | 1.13 ± 1.03 | 0.55 ± 0.59 | 40 |
| ***plt2, plt3*** | 1.26 ± 1.06 | 0.55 ± 0.74 | 65 |
| ***plt3 pPLT3::PLT3-mV*** | 0.95 ± 1.01 | 0.88 ± 0.62 | 104 |
| ***plt2, plt3 pPLT3::PLT3-mV*** | 1.40 ± 1.05 | 0.66 ± 0.62 | 62 |
| ***plt2, plt3 pPLT3::*** **PLT3ΔPrD-mV** | 1.34 ± 1.09 | 0.48 ± 0.62 | 44 |

**Supplementary Table 10: FLIM results of subnuclear data analysis in WOX5-mV and PLT3-mCh co-expressing *N. benthamiana* epidermal cells related to Figure 5.**

| date | measurement # | WOX5-mVenus fluorescence lifetime [ns] | |
|---|---|---|---|
| | | bodies-only | nucleoplasm |
| 2019-06-11 | WOX5-mV PLT3-mCh_7 | 2.24 | 2.76 |
| | WOX5-mV PLT3-mCh_15 | 2.19 | 2.85 |
| 2019-06-10 | WOX5-mV PLT3-mCh_23 | 2.60 | 2.92 |
| | WOX5-mV PLT3-mCh_25 | 2.55 | 2.88 |
| 2018-06-29 | WOX5-mV PLT3-mCh_4 | 2.56 | 2.89 |
| 2018-06-28 | WOX5-mV PLT3-mCh_1 | 2.50 | 2.84 |
| | WOX5-mV PLT3-mCh_9 | 2.50 | 2.69 |
| **mean** | | **2.45** | **2.83** |
| **st. dev.** | | **0.15** | **0.07** |

### Supplementary Figure 11 - SCN raw data

| date | root # | *Col* | | *plt3-1* | | *plt2* | | *plt2, plt3* | | *wox5-1* | | *wox5, plt3* | | *wox5, plt2, plt3* | |
|---|---|---|---|---|---|---|---|---|---|---|---|---|---|---|---|
| 2017-11-23 | | QC divisions | CSC layers | QC divisions | CSC layers | QC divisions | CSC layers | QC divisions | CSC layers | QC divisions | CSC layers | QC divisions | CSC layers | QC divisions | CSC layers |
| | 1 | 2 | 1 | 2 | 2 | 1 | 1 | 0 | 1 | 2 | 0 | 4 | 0 | | |
| | 2 | 0 | 1 | 2 | 2 | 0 | 1 | 1 | 2 | 4 | 0 | 3 | 0 | | |
| | 3 | 0 | 2 | 1 | 1 | 0 | 1 | 1 | 2 | 1 | 0 | 2 | 0 | | |
| | 4 | 0 | 1 | 0 | 1 | 1 | 2 | 0 | 1 | 3 | 1 | 1 | 1 | | |
| | 5 | 1 | 1 | 0 | 1 | 1 | 1 | 0 | 1 | 0 | 0 | 2 | 1 | | |
| | 6 | 1 | 1 | 0 | 1 | 1 | 1 | 1 | 2 | 2 | 0 | 1 | 0 | | |
| | 7 | 0 | 1 | 0 | 1 | 0 | 0 | 2 | 3 | 2 | 1 | 2 | 2 | | |
| | 8 | 0 | 2 | 0 | 1 | 1 | 2 | 0 | 1 | 2 | 0 | 0 | 0 | | |
| | 9 | 0 | 0 | 0 | 1 | 0 | 1 | 0 | 1 | 2 | 0 | 2 | 1 | | |
| | 10 | 1 | 2 | 0 | 0 | 0 | 2 | 0 | 1 | 1 | 0 | 1 | 1 | | |
| | 11 | 1 | 2 | 1 | 2 | 2 | 1 | 1 | 1 | 2 | 0 | 2 | 1 | | |
| | 12 | 0 | 1 | 0 | 0 | 1 | 1 | 0 | 2 | 2 | 0 | 2 | 0 | | |
| | 13 | 0 | 2 | 0 | 0 | 0 | 0 | 0 | 0 | 2 | 0 | 2 | 0 | | |
| | 14 | 0 | 2 | 0 | 1 | 1 | 2 | 0 | 2 | 1 | 0 | 1 | 0 | | |
| | 15 | 0 | 1 | 0 | 1 | 1 | 1 | 2 | 1 | 0 | 0 | 3 | 0 | | |
| | 16 | 0 | 2 | 1 | 0 | 1 | 0 | 1 | 0 | 2 | 0 | 1 | 0 | | |
| | 17 | 1 | 1 | 1 | 1 | 0 | 2 | 0 | 2 | 2 | 0 | 1 | 1 | | |
| | 18 | 0 | 0 | 0 | 1 | 0 | 1 | 0 | 0 | 1 | 0 | 2 | 0 | | |
| | 19 | 0 | 2 | 0 | 1 | 0 | 0 | 1 | 2 | 1 | 1 | 1 | 0 | | |
| | 20 | 0 | 1 | 1 | 0 | 0 | 1 | 0 | 0 | 1 | 0 | 2 | 0 | | |
| | 21 | 0 | 1 | 0 | 0 | 0 | 0 | 1 | 1 | 2 | 0 | 1 | 0 | | |
| | 22 | | | 0 | 1 | 1 | 0 | 1 | 1 | 2 | 1 | 1 | 0 | | |
| | 23 | | | 0 | 0 | 0 | 1 | 0 | 1 | 1 | 1 | 3 | 0 | | |
| | 24 | | | 0 | 2 | 0 | 1 | 2 | 0 | 1 | 0 | 1 | 0 | | |
| | 25 | | | | | 0 | 0 | 1 | 2 | 1 | 0 | 1 | 0 | | |
| | 26 | | | | | 0 | 1 | 0 | 1 | 2 | 0 | 2 | 0 | | |
| | 27 | | | | | 0 | 2 | 1 | 0 | 0 | 0 | | | | |
| | 28 | | | | | 2 | 1 | | | 1 | 0 | | | | |
| | 29 | | | | | 2 | 0 | | | | | | | | |
| | 30 | | | | | 1 | 1 | | | | | | | | |
| | 31 | | | | | 0 | 1 | | | | | | | | |
| | 32 | | | | | 0 | 1 | | | | | | | | |
| 2017-12-05 | | QC divisions | CSC layers | QC divisions | CSC layers | QC divisions | CSC layers | QC divisions | CSC layers | QC divisions | CSC layers | QC divisions | CSC layers | QC divisions | CSC layers |
| | 1 | 0 | 2 | 1 | 0 | | | 0 | 1 | 2 | 1 | 2 | 2 | | |
| | 2 | 0 | 1 | 1 | 2 | | | 2 | 0 | 2 | 0 | 4 | 0 | | |
| | 3 | 0 | 1 | 0 | 2 | | | 1 | 0 | 3 | 0 | 2 | 0 | | |
| | 4 | 0 | 0 | 1 | 2 | | | 0 | 2 | 3 | 0 | 1 | 1 | | |
| | 5 | 0 | 1 | 1 | 1 | | | 0 | 2 | 2 | 0 | 2 | 1 | | |
| | 6 | 1 | 0 | 2 | 1 | | | 0 | 1 | 1 | 0 | 2 | 0 | | |
| | 7 | 0 | 1 | 1 | 1 | | | 0 | 1 | 2 | 0 | 2 | 1 | | |
| | 8 | 0 | 1 | 0 | 1 | | | 2 | 1 | 2 | 0 | 3 | 1 | | |
| | 9 | 0 | 1 | 0 | 0 | | | 1 | 0 | 3 | 0 | 1 | 0 | | |
| | 10 | 0 | 1 | 2 | 2 | | | 0 | 2 | 2 | 0 | 1 | 1 | | |
| | 11 | 0 | 1 | 1 | 2 | | | 2 | 1 | 2 | 1 | 1 | 0 | | |
| | 12 | 0 | 1 | 0 | 2 | | | 0 | 2 | 2 | 0 | 4 | 1 | | |
| | 13 | 0 | 1 | 1 | 0 | | | 0 | 1 | 0 | 1 | 1 | 0 | | |
| | 14 | 1 | 1 | 1 | 1 | | | 2 | 0 | 1 | 0 | 2 | 1 | | |
| | 15 | 0 | 0 | 0 | 1 | | | 1 | 0 | 1 | 0 | 2 | 0 | | |
| | 16 | 0 | 1 | 0 | 1 | | | 1 | 0 | 1 | 0 | 2 | 1 | | |
| | 17 | 0 | 1 | 0 | 2 | | | 0 | 1 | 3 | 0 | 4 | 0 | | |
| | 18 | 0 | 0 | 0 | 1 | | | 0 | 1 | 3 | 0 | 3 | 1 | | |
| | 19 | 0 | 1 | 1 | 1 | | | 1 | 0 | 2 | 1 | 2 | 0 | | |
| | 20 | 0 | 2 | 1 | 2 | | | 1 | 1 | 1 | 0 | 2 | 0 | | |
| | 21 | 0 | 0 | 0 | 1 | | | 3 | 0 | 2 | 1 | 3 | 0 | | |
| | 22 | 0 | 1 | 0 | 0 | | | 1 | 0 | 2 | 0 | 3 | 1 | | |
| | 23 | 0 | 1 | 1 | 1 | | | | | 3 | 0 | 3 | 0 | | |
| | 24 | 2 | 1 | 0 | 0 | | | | | 3 | 0 | 3 | 0 | | |
| | 25 | 0 | 1 | 0 | 1 | | | | | 2 | 1 | 3 | 0 | | |
| | 26 | | | 0 | 2 | | | | | 1 | 0 | 2 | 1 | | |
| | 27 | | | 2 | 0 | | | | | 3 | 0 | | | | |
| | 28 | | | 1 | 0 | | | | | 1 | 1 | | | | |
| 2018-05-01 | | QC divisions | CSC layers | QC divisions | CSC layers | QC divisions | CSC layers | QC divisions | CSC layers | QC divisions | CSC layers | QC divisions | CSC layers | QC divisions | CSC layers |
| | 1 | 1 | 0 | 1 | 2 | 0 | 1 | 1 | 1 | 3 | 0 | 2 | 0 | | |
| | 2 | 0 | 1 | 1 | 0 | 1 | 2 | 0 | 2 | 1 | 0 | 3 | 0 | | |
| | 3 | 0 | 0 | 1 | 1 | 0 | 0 | 0 | 0 | 3 | 0 | 1 | 0 | | |
| | 4 | 1 | 1 | 0 | 1 | 0 | 1 | 0 | 0 | 1 | 0 | 3 | 1 | | |
| | 5 | 0 | 1 | 0 | 2 | 1 | 1 | 3 | 0 | 3 | 1 | 1 | 0 | | |
| | 6 | 0 | 1 | 0 | 1 | 0 | 0 | 1 | 2 | 2 | 0 | 4 | 0 | | |
| | 7 | 0 | 2 | 1 | 1 | 1 | 1 | 2 | 2 | 2 | 0 | 2 | 1 | | |
| | 8 | 2 | 1 | 0 | 1 | 1 | 1 | 2 | 0 | 3 | 0 | 0 | 0 | | |
| | 9 | 1 | 1 | 0 | 2 | 1 | 1 | 0 | 1 | 0 | 0 | 3 | 0 | | |
| | 10 | 1 | 0 | 1 | 0 | 1 | 1 | 1 | 1 | 1 | 0 | 3 | 0 | | |
| | 11 | 1 | 1 | 0 | 0 | 2 | 0 | 1 | 1 | 3 | 0 | 2 | 0 | | |
| | 12 | 1 | 0 | 0 | 1 | 0 | 2 | 0 | 0 | 2 | 0 | 3 | 0 | | |
| | 13 | 0 | 1 | 0 | 1 | 2 | 2 | 1 | 1 | 1 | 1 | 1 | 1 | | |
| | 14 | 1 | 2 | 1 | 1 | 1 | 2 | 0 | 1 | 3 | 0 | 3 | 0 | | |
| | 15 | 1 | 1 | 1 | 1 | 0 | 0 | 0 | 2 | 2 | 0 | 4 | 0 | | |
| | 16 | 1 | 1 | 1 | 1 | 0 | 0 | 0 | 0 | 1 | 1 | 2 | 0 | | |
| | 17 | 2 | 1 | 2 | 1 | 1 | 0 | 1 | 1 | 3 | 0 | 2 | 0 | | |
| | 18 | 0 | 1 | 2 | 1 | 1 | 1 | 1 | 1 | 2 | 0 | 1 | 0 | | |
| | 19 | 1 | 1 | 2 | 1 | 1 | 0 | 0 | 1 | 3 | 0 | 3 | 0 | | |
| | 20 | 1 | 1 | 2 | 0 | 0 | 1 | 1 | 1 | 3 | 0 | 3 | 0 | | |
| | 21 | 0 | 1 | 0 | 1 | 1 | 1 | 1 | 0 | 2 | 0 | 2 | 0 | | |
| | 22 | 2 | 1 | 0 | 1 | 1 | 2 | 0 | 0 | 1 | 1 | 2 | 0 | | |
| | 23 | 0 | 1 | 0 | 2 | 0 | 1 | 0 | 1 | 4 | 0 | 3 | 0 | | |
| | 24 | 1 | 0 | 0 | 2 | 0 | 1 | 1 | 0 | 3 | 0 | 3 | 1 | | |
| | 25 | 0 | 1 | 0 | 2 | 1 | 0 | 1 | 0 | 3 | 0 | 3 | 0 | | |
| | 26 | 0 | 0 | 1 | 1 | 0 | 0 | 0 | 0 | 3 | 0 | 3 | 0 | | |
| | 27 | 0 | 1 | 2 | 0 | 2 | 0 | 1 | 0 | 3 | 0 | 1 | 0 | | |
| | 28 | 0 | 1 | 1 | 1 | 0 | 0 | 1 | 0 | 1 | 0 | 1 | 1 | | |
| | 29 | 0 | 1 | 0 | 2 | 1 | 0 | 1 | 1 | 1 | 0 | 1 | 1 | | |
| | 30 | 0 | 1 | 2 | 1 | 0 | 0 | 2 | 0 | 2 | 0 | 4 | 0 | | |
| | 31 | 0 | 2 | 0 | 1 | 0 | 0 | 0 | 0 | 2 | 1 | 3 | 0 | | |
| | 32 | 0 | 1 | 0 | 1 | 0 | 0 | 0 | 2 | 3 | 0 | 1 | 0 | | |
| | 33 | 0 | 0 | 1 | 1 | 1 | 1 | 1 | 0 | | | 3 | 0 | | |
| | 34 | 0 | 1 | 2 | 1 | 0 | 0 | | | | | 2 | 0 | | |
| | 35 | 1 | 1 | 1 | 1 | 0 | 2 | | | | | 2 | 0 | | |
| | 36 | 0 | 1 | 0 | 1 | 0 | 1 | | | | | | | | |
| | 37 | | | 0 | 1 | 1 | 0 | | | | | | | | |
| | 38 | | | 1 | 1 | 0 | 0 | | | | | | | | |
| | 39 | | | 1 | 0 | 0 | 1 | | | | | | | | |
| | 40 | | | 0 | 0 | 1 | 2 | | | | | | | | |
| | 41 | | | 1 | 1 | 0 | 1 | | | | | | | | |
| | 42 | | | 2 | 0 | 0 | 2 | | | | | | | | |
| | 43 | | | 1 | 1 | 1 | 0 | | | | | | | | |
| | 44 | | | 0 | 1 | 0 | 0 | | | | | | | | |
| | 45 | | | 1 | 0 | 0 | 1 | | | | | | | | |
| | 46 | | | 0 | 1 | | | | | | | | | | |
| | 47 | | | 1 | 0 | | | | | | | | | | |
| | 48 | | | 0 | 2 | | | | | | | | | | |
| | 49 | | | 0 | 0 | | | | | | | | | | |
| | 50 | | | 0 | 1 | | | | | | | | | | |
| | 51 | | | 1 | 0 | | | | | | | | | | |
| 2018-08-01 | | QC divisions | CSC layers | QC divisions | CSC layers | QC divisions | CSC layers | QC divisions | CSC layers | QC divisions | CSC layers | QC divisions | CSC layers | QC divisions | CSC layers |

### Supplementary Figure 11 - SCN raw data

| | | | | | | | | | | | | | | | |
|---|---|---|---|---|---|---|---|---|---|---|---|---|---|---|---|
| | 1 | | | | | | | | | | | | | 4 | 0 |
| | 2 | | | | | | | | | | | | | 2 | 0 |
| | 3 | | | | | | | | | | | | | 2 | 0 |
| | 4 | | | | | | | | | | | | | 3 | 0 |
| | 5 | | | | | | | | | | | | | 4 | 1 |
| | 6 | | | | | | | | | | | | | 3 | 0 |
| | 7 | | | | | | | | | | | | | 2 | 1 |
| | 8 | | | | | | | | | | | | | 2 | 0 |
| | 9 | | | | | | | | | | | | | 3 | 0 |
| | 10 | | | | | | | | | | | | | 3 | 0 |
| | 11 | | | | | | | | | | | | | 2 | 0 |
| | 12 | | | | | | | | | | | | | 2 | 1 |
| | 13 | | | | | | | | | | | | | 4 | 0 |
| | 14 | | | | | | | | | | | | | 3 | 0 |
| | 15 | | | | | | | | | | | | | 4 | 0 |
| | 16 | | | | | | | | | | | | | 4 | 0 |
| | 17 | | | | | | | | | | | | | 3 | 0 |
| | 18 | | | | | | | | | | | | | 4 | 0 |
| | 19 | | | | | | | | | | | | | 4 | 0 |
| | 20 | | | | | | | | | | | | | 4 | 1 |
| | 21 | | | | | | | | | | | | | 4 | 0 |
| | 22 | | | | | | | | | | | | | 2 | 0 |
| | 23 | | | | | | | | | | | | | 3 | 0 |
| | 24 | | | | | | | | | | | | | 4 | 0 |
| | 25 | | | | | | | | | | | | | 4 | 1 |
| | 26 | | | | | | | | | | | | | 4 | 1 |
| | 27 | | | | | | | | | | | | | 3 | 0 |
| | 28 | | | | | | | | | | | | | 0 | 1 |
| | 29 | | | | | | | | | | | | | 3 | 0 |
| | 30 | | | | | | | | | | | | | 2 | 1 |
| | 31 | | | | | | | | | | | | | 1 | 0 |
| | 32 | | | | | | | | | | | | | 4 | 0 |
| | 33 | | | | | | | | | | | | | 4 | 0 |
| | 34 | | | | | | | | | | | | | 2 | 0 |
| | 35 | | | | | | | | | | | | | 3 | 0 |
| | 36 | | | | | | | | | | | | | 3 | 0 |
| **2018-08-30** | | **QC divisions** | **CSC layers** | **QC divisions** | **CSC layers** | **QC divisions** | **CSC layers** | **QC divisions** | **CSC layers** | **QC divisions** | **CSC layers** | **QC divisions** | **CSC layers** | **QC divisions** | **CSC layers** |
| | 1 | | | | | | | | | | | | | 3 | 1 |
| | 2 | | | | | | | | | | | | | 4 | 0 |
| | 3 | | | | | | | | | | | | | 2 | 0 |
| | 4 | | | | | | | | | | | | | 3 | 0 |
| | 5 | | | | | | | | | | | | | 4 | 0 |
| | 6 | | | | | | | | | | | | | 3 | 0 |
| | 7 | | | | | | | | | | | | | 4 | 1 |
| | 8 | | | | | | | | | | | | | 2 | 0 |
| | 9 | | | | | | | | | | | | | 2 | 0 |
| | 10 | | | | | | | | | | | | | 3 | 1 |
| | 11 | | | | | | | | | | | | | 3 | 0 |
| | 12 | | | | | | | | | | | | | 1 | 0 |
| | 13 | | | | | | | | | | | | | 3 | 0 |
| | 14 | | | | | | | | | | | | | 4 | 1 |
| | 15 | | | | | | | | | | | | | 4 | 1 |
| | 16 | | | | | | | | | | | | | 3 | 1 |
| | 17 | | | | | | | | | | | | | 4 | 0 |
| | 18 | | | | | | | | | | | | | 1 | 0 |
| | 19 | | | | | | | | | | | | | 2 | 0 |
| | 20 | | | | | | | | | | | | | 2 | 0 |
| | 21 | | | | | | | | | | | | | 4 | 1 |
| | 22 | | | | | | | | | | | | | 4 | 0 |
| | 23 | | | | | | | | | | | | | 4 | 0 |
| | 24 | | | | | | | | | | | | | 2 | 0 |
| | 25 | | | | | | | | | | | | | 1 | 0 |
| | 26 | | | | | | | | | | | | | 2 | 0 |
| **2019-04-01** | | **QC divisions** | **CSC layers** | **QC divisions** | **CSC layers** | **QC divisions** | **CSC layers** | **QC divisions** | **CSC layers** | **QC divisions** | **CSC layers** | **QC divisions** | **CSC layers** | **QC divisions** | **CSC layers** |
| | 1 | 1 | 2 | | | | | | | | | | | 1 | 0 |
| | 2 | 0 | 1 | | | | | | | | | | | 1 | 0 |
| | 3 | 0 | 1 | | | | | | | | | | | 4 | 0 |
| | 4 | 0 | 1 | | | | | | | | | | | 2 | 1 |
| | 5 | 0 | 1 | | | | | | | | | | | 4 | 0 |
| | 6 | 0 | 2 | | | | | | | | | | | 3 | 0 |
| | 7 | 0 | 1 | | | | | | | | | | | 3 | 0 |
| | 8 | 0 | 1 | | | | | | | | | | | 4 | 0 |
| | 9 | 2 | 0 | | | | | | | | | | | 3 | 1 |
| | 10 | 0 | 1 | | | | | | | | | | | 3 | 0 |
| | 11 | 0 | 1 | | | | | | | | | | | 1 | 2 |
| | 12 | 0 | 0 | | | | | | | | | | | 3 | 0 |
| | 13 | 0 | 1 | | | | | | | | | | | 2 | 0 |
| | 14 | 0 | 1 | | | | | | | | | | | 2 | 0 |
| | 15 | 1 | 0 | | | | | | | | | | | 1 | 0 |
| | 16 | 0 | 1 | | | | | | | | | | | 3 | 0 |
| | 17 | 0 | 1 | | | | | | | | | | | 3 | 1 |
| | 18 | 0 | 1 | | | | | | | | | | | 4 | 0 |
| | 19 | 1 | 1 | | | | | | | | | | | 3 | 1 |
| | 20 | 0 | 2 | | | | | | | | | | | 4 | 1 |
| | 21 | 0 | 1 | | | | | | | | | | | 4 | 0 |
| | 22 | 0 | 1 | | | | | | | | | | | 4 | 0 |
| | 23 | 0 | 2 | | | | | | | | | | | 3 | 0 |
| | 24 | 0 | 3 | | | | | | | | | | | 3 | 0 |
| | 25 | 1 | 1 | | | | | | | | | | | 2 | 0 |
| | 26 | 0 | 1 | | | | | | | | | | | 2 | 0 |
| | 27 | 0 | 1 | | | | | | | | | | | 2 | 0 |
| | 28 | 2 | 1 | | | | | | | | | | | 3 | 1 |
| | 29 | 0 | 2 | | | | | | | | | | | 3 | 1 |
| | 30 | 0 | 1 | | | | | | | | | | | 2 | 0 |
| | 31 | 0 | 1 | | | | | | | | | | | 3 | 0 |
| | 32 | 0 | 2 | | | | | | | | | | | 0 | 0 |
| | 33 | 1 | 1 | | | | | | | | | | | 3 | 0 |
| | 34 | 0 | 1 | | | | | | | | | | | 2 | 0 |
| | 35 | | | | | | | | | | | | | 4 | 0 |
| | 36 | | | | | | | | | | | | | 1 | 0 |
| **2019-05-01** | | **QC divisions** | **CSC layers** | **QC divisions** | **CSC layers** | **QC divisions** | **CSC layers** | **QC divisions** | **CSC layers** | **QC divisions** | **CSC layers** | **QC divisions** | **CSC layers** | **QC divisions** | **CSC layers** |
| | 1 | 0 | 1 | 1 | 0 | | | 0 | 1 | 2 | 0 | | | | |
| | 2 | 0 | 1 | 0 | 0 | | | 0 | 1 | 2 | 0 | | | | |
| | 3 | 0 | 1 | 1 | 0 | | | 1 | 0 | 4 | 0 | | | | |
| | 4 | 0 | 0 | 0 | 0 | | | 1 | 0 | 2 | 0 | | | | |
| | 5 | 0 | 1 | 2 | 0 | | | 1 | 3 | 3 | 0 | | | | |
| | 6 | 0 | 2 | 0 | 0 | | | 0 | 2 | 4 | 0 | | | | |
| | 7 | 0 | 1 | 0 | 1 | | | 2 | 2 | 0 | 1 | | | | |
| | 8 | 0 | 1 | 0 | 1 | | | 0 | 1 | 2 | 0 | | | | |
| | 9 | 0 | 1 | 1 | 0 | | | 4 | 2 | 1 | 0 | | | | |
| | 10 | 0 | 1 | 1 | 0 | | | 2 | 0 | 1 | 0 | | | | |
| | 11 | 0 | 0 | 1 | 1 | | | 0 | 0 | 3 | 1 | | | | |
| | 12 | 0 | 1 | 1 | 0 | | | 0 | 0 | 1 | 1 | | | | |
| | 13 | 1 | 1 | 1 | 1 | | | 1 | 0 | 1 | 0 | | | | |
| | 14 | 0 | 1 | 0 | 0 | | | 3 | 0 | 2 | 0 | | | | |
| | 15 | 0 | 0 | 0 | 0 | | | 2 | 0 | 3 | 0 | | | | |

### Supplementary Figure 11 - SCN raw data

| | | | | | | | | | | | | | | |
|---|---|---|---|---|---|---|---|---|---|---|---|---|---|---|
| 16 | 0 | 1 | 1 | 0 | | | 1 | 1 | 1 | 1 | | | | |
| 17 | 0 | 1 | 1 | 0 | | | 0 | 0 | 0 | 1 | | | | |
| 18 | 1 | 0 | 2 | 0 | | | 1 | 0 | 1 | 0 | | | | |
| 19 | 0 | 2 | 0 | 1 | | | 1 | 1 | 3 | 0 | | | | |
| 20 | 0 | 1 | 0 | 0 | | | 3 | 2 | 1 | 2 | | | | |
| 21 | 1 | 1 | | | | | 1 | 2 | 1 | 0 | | | | |
| 22 | 1 | 0 | | | | | 2 | 0 | 0 | 0 | | | | |
| 23 | 1 | 1 | | | | | 1 | 0 | 0 | 1 | | | | |
| 24 | 0 | 1 | | | | | | | 2 | 0 | | | | |
| 25 | 0 | 0 | | | | | | | 3 | 0 | | | | |
| 26 | 2 | 0 | | | | | | | 2 | 0 | | | | |
| 27 | 2 | 0 | | | | | | | 2 | 1 | | | | |
| 28 | 0 | 1 | | | | | | | 1 | 1 | | | | |
| 29 | 0 | 1 | | | | | | | 2 | 0 | | | | |
| 30 | 1 | 2 | | | | | | | 3 | 0 | | | | |
| mean | 0.34 | 0.99 | 0.61 | 0.84 | 0.53 | 0.82 | 0.81 | 0.84 | 1.87 | 0.23 | 2.11 | 0.31 | 2.83 | 0.24 |
| SD | 0.59 | 0.59 | 0.69 | 0.69 | 0.64 | 0.72 | 0.87 | 0.82 | 0.98 | 0.44 | 0.98 | 0.51 | 1.04 | 0.45 |
| n | 146 | 146 | 123 | 123 | 77 | 77 | 105 | 105 | 118 | 118 | 87 | 87 | 98 | 98 |

### Supplementary Table 12 - FLIM raw data

| date | measurement # | mVenus fluorescence lifetime [ns] | | | | | | | |
|---|---|---|---|---|---|---|---|---|---|
| | | WOX5-mV | + free mCh | + PLT1-mCh | + PLT2-mCh | + PLT3-mCh | + PLT4-mCh | + PLT3ΔQ-mCh | + PLT3ΔPrD-mCh |
| 2018-06-28 | 1 | 3.032 | | | | 2.84 | | | |
| | 2 | 3.01 | | | | 2.959 | | | |
| | 3 | 2.95 | | | | 2.682 | | | |
| | 4 | 3.033 | | | | 2.81 | | | |
| | 5 | 3 | | | | 2.802 | | | |
| | 6 | 2.991 | | | | 2.7888 | | | |
| | 7 | 3.033 | | | | 2.58 | | | |
| | 8 | 3.038 | | | | 2.55 | | | |
| | 9 | 3 | | | | 2.58 | | | |
| | 10 | 3.004 | | | | 2.74 | | | |
| 2018-06-29 | 1 | 3.07 | 3.06 | 2.96503 | 2.85 | 2.97589 | | | |
| | 2 | 3.052 | 3.02955 | 2.86599 | 2.80095 | 2.85021 | | | |
| | 3 | 3.1 | 2.76226 | 2.75013 | 2.72653 | 2.2656 | | | |
| | 4 | 3.059 | 3.06 | 2.83846 | 2.72653 | 2.83412 | | | |
| | 5 | 3.065 | 3.00629 | 2.57539 | 2.97515 | 2.43652 | | | |
| | 6 | 3.03 | 2.98834 | 2.79689 | 2.81017 | 2.93302 | | | |
| | 7 | 3.052 | 2.98048 | 2.87333 | 2.7397 | 2.77297 | | | |
| | 8 | 3.065 | 2.95994 | 2.90885 | 2.77644 | 2.69642 | | | |
| | 9 | 3.073 | 3.01475 | 2.92286 | 2.6488 | 2.55564 | | | |
| | 10 | 3.053 | 3.02998 | 2.95125 | 2.72693 | 2.76554 | | | |
| | 11 | | | 2.7025 | 2.46182 | | | | |
| | 12 | | | 2.84014 | 2.87404 | | | | |
| | 13 | | | 2.88 | 2.85923 | | | | |
| | 14 | | | 2.87274 | 2.70974 | | | | |
| | 15 | | | 2.84029 | 2.60848 | | | | |
| | 16 | | | 2.80994 | 2.39638 | | | | |
| | 17 | | | 2.72154 | 2.52347 | | | | |
| | 18 | | | 2.85102 | 2.92471 | | | | |
| | 19 | | | 2.63374 | 2.7354 | | | | |
| | 20 | | | 2.60274 | 2.67349 | | | | |
| | 21 | | | | 2.73458 | | | | |
| 2018-07-04 | 1 | 2.994 | 2.956 | 2.912 | 2.76679 | 2.91558 | | | |
| | 2 | 2.96 | 2.952 | 2.57812 | 2.83663 | 2.50295 | | | |
| | 3 | 2.982 | 2.938 | 2.63027 | 2.59163 | 2.6749 | | | |
| | 4 | 2.98 | 2.88501 | 2.77968 | 2.73952 | 2.63889 | | | |
| | 5 | 2.995 | 2.97083 | 2.6332 | 2.6176 | 2.58327 | | | |
| | 6 | 2.998 | 2.92704 | 2.963 | 2.60066 | 2.65011 | | | |
| | 7 | 3.022 | 2.87015 | 2.72182 | 2.76336 | 2.5485 | | | |
| | 8 | 2.98 | 2.8572 | 2.81864 | 2.84407 | 2.72492 | | | |
| | 9 | 2.987 | 2.96015 | 2.80822 | 2.78187 | 2.76272 | | | |
| | 10 | 2.966 | 2.95641 | 2.70145 | 2.74231 | 2.57421 | | | |
| | 11 | | | 2.87194 | 2.67259 | | | | |
| | 12 | | | 2.75764 | 2.68184 | | | | |
| | 13 | | | 2.8172 | 2.62721 | | | | |
| | 14 | | | 2.78794 | 2.80769 | | | | |
| | 15 | | | 2.58113 | 2.94565 | | | | |
| 2019-05-13 | 1 | 3.039 | 3.036 | | | 2.962 | | 2.986 | 3 |
| | 2 | 3.055 | 3.033 | | | 2.977 | | 2.97 | 3.03 |
| | 3 | 3.03 | 3.032 | | | 2.861 | | 2.891 | 3.02 |
| | 4 | 3.032 | 2.969 | | | 2.92 | | 2.76 | 3.038 |
| | 5 | 3.014 | 2.899 | | | 3.023 | | 2.818 | 2.89 |
| | 6 | 2.995 | 2.875 | | | 2.98 | | 2.73 | 2.76 |
| | 7 | 3.01 | 3.034 | | | 2.92 | | 2.81 | 2.67 |
| | 8 | 3.01 | 2.977 | | | 2.874 | | 3.037 | 2.47 |
| | 9 | 3.071 | 3.004 | | | 2.727 | | 2.94 | 2.653 |
| | 10 | 3.049 | 2.92 | | | 2.77 | | 3.01 | 2.95 |
| | 11 | | | | | 2.711 | | 2.978 | 2.967 |
| | 12 | | | | | 2.758 | | 2.969 | 2.91 |
| | 13 | | | | | | | 2.74 | 2.8 |
| | 14 | | | | | | | 2.997 | 2.92 |
| | 15 | | | | | | | 2.93 | 2.96 |
| | 16 | | | | | | | 2.736 | 2.95 |
| | 17 | | | | | | | 2.712 | 2.72 |
| | 18 | | | | | | | 2.86 | 2.97 |
| | 19 | | | | | | | 2.956 | 2.872 |
| | 20 | | | | | | | 2.895 | 2.79 |
| 2019-05-14 | 1 | 3.03 | 3.04 | | | | | 3.123 | 2.9852 |
| | 2 | 3.02 | 3.039 | | | | | 3.034 | 2.904 |
| | 3 | 3.045 | 3.049 | | | | | 2.96 | 2.937 |
| | 4 | 3.076 | 2.833 | | | | | 2.983 | 2.957 |
| | 5 | 3.058 | 3 | | | | | 3.06 | 2.846 |
| | 6 | 3.076 | 3.06 | | | | | 3.073 | 2.92 |
| | 7 | 3.046 | 3.047 | | | | | 3.119 | 2.88 |
| | 8 | 3.033 | | | | | | 3.079 | 2.989 |
| | 9 | 3.028 | | | | | | 3.074 | 2.974 |
| | 10 | 3.065 | | | | | | 3.048 | 2.894 |
| | 11 | 3.067 | | | | | | | 2.92 |
| | 12 | 3.051 | | | | | | | 2.709 |
| 2015-10-27 | 1 | 3.097 | | | | | | 3.017 | |
| | 2 | 3.084 | | | | | | 3.116 | |
| | 3 | 3.083 | | | | | | 3.01 | |
| | 4 | 3.065 | | | | | | 2.62 | |
| | 5 | 3.036 | | | | | | 2.931 | |
| | 6 | 3.073 | | | | | | 2.937 | |
| | 7 | 3.055 | | | | | | 2.942 | |
| | 8 | 3.063 | | | | | | 2.805 | |
| | 9 | 3.033 | | | | | | 3.17 | |
| | 10 | 3.061 | | | | | | 2.457 | |

### Supplementary Table 12 - FLIM raw data

| | | | | | | | | | |
|---|---|---|---|---|---|---|---|---|---|
| | 11 | | | | | | | 3.11 | |
| | 12 | | | | | | | 3.02 | |
| | 13 | | | | | | | 3.017 | |
| | 14 | | | | | | | 3.029 | |
| | 15 | | | | | | | 3.02 | |
| | 16 | | | | | | | 2.99 | |
| | 17 | | | | | | | 2.995 | |
| | 18 | | | | | | | 2.95 | |
| | 19 | | | | | | | 2.943 | |
| | 20 | | | | | | | 2.77 | |
| **2021-02-05** | **1** | 3.124 | 3.01 | | | | 2.774 | | |
| | **2** | 3.08 | 3.005 | | | | 2.771 | | |
| | **3** | 3.135 | 3.037 | | | | 2.53 | | |
| | **4** | 3.1 | 3.01 | | | | 2.717 | | |
| | **5** | 3.127 | 3.025 | | | | 2.44 | | |
| | **6** | 3.142 | 3.038 | | | | 2.57 | | |
| | **7** | 3.14 | 3.041 | | | | 2.374 | | |
| | **8** | 3.168 | 3 | | | | 2.56 | | |
| | **9** | 3.119 | 3.025 | | | | 2.551 | | |
| | **10** | 3.141 | 3.031 | | | | 2.588 | | |
| | **11** | 3.108 | 2.996 | | | | 2.8 | | |
| | **12** | 3.169 | 2.96 | | | | 2.67 | | |
| | **13** | | 2.97 | | | | 2.557 | | |
| | **14** | | 2.97 | | | | 2.555 | | |
| | **15** | | 3.024 | | | | 2.59 | | |
| | **16** | | 3 | | | | 2.583 | | |
| | **17** | | 2.93 | | | | 1.819 | | |
| | **18** | | 2.95 | | | | 2.857 | | |
| | **19** | | 3.005 | | | | 2.8 | | |
| | **20** | | 3 | | | | 2.18 | | |
| **2021-03-17** | **1** | 2.975 | | | | | 2.678 | | |
| | **2** | 2.93 | | | | | 2.588 | | |
| | **3** | 2.992 | | | | | 2.966 | | |
| | **4** | 2.964 | | | | | 2.901 | | |
| | **5** | 2.832 | | | | | 2.717 | | |
| | **6** | 2.91 | | | | | 2.257 | | |
| | **7** | 2.79 | | | | | 2.23 | | |
| | **8** | 2.94 | | | | | 2.99 | | |
| | **9** | 2.893 | | | | | 3.011 | | |
| | **10** | 2.964 | | | | | 2.955 | | |
| | **11** | | | | | | 2.89 | | |
| | **12** | | | | | | 2.75 | | |
| | **13** | | | | | | 2.803 | | |
| | **14** | | | | | | 2.7 | | |
| | **15** | | | | | | 2.803 | | |
| | **16** | | | | | | 2.74 | | |
| | **17** | | | | | | 2.74 | | |
| | **18** | | | | | | 3.001 | | |
| | **19** | | | | | | 2.97 | | |
| | **20** | | | | | | 2.8 | | |
| **2021-03-30** | **1** | 2.77 | | | | | 3.01 | | |
| | **2** | 2.99 | | | | | 3.056 | | |
| | **3** | 3.058 | | | | | 2.977 | | |
| | **4** | 3.037 | | | | | 2.935 | | |
| | **5** | 3.08 | | | | | 2.761 | | |
| | **6** | 3.076 | | | | | 2.69 | | |
| | **7** | 3.073 | | | | | 2.85 | | |
| | **8** | 3.04 | | | | | 3.082 | | |
| | **9** | 3 | | | | | 3.027 | | |
| | **10** | 3.094 | | | | | 3.03 | | |
| | **11** | | | | | | 2.973 | | |
| | **12** | | | | | | 2.91 | | |
| | **13** | | | | | | 2.905 | | |
| | **14** | | | | | | 2.912 | | |
| | **15** | | | | | | 2.906 | | |
| | **16** | | | | | | 2.882 | | |
| | **17** | | | | | | 2.84 | | |
| | **18** | | | | | | 2.884 | | |
| | **19** | | | | | | 2.9 | | |
| | **20** | | | | | | 2.742 | | |
| | **n** | **94** | **57** | **35** | **36** | **42** | **60** | **50** | **32** |
| | **mean** | **3.03** | **2.98** | **2.79** | **2.73** | **2.75** | **2.75** | **2.94** | **2.88** |
| | **SD** | **0.07** | **0.06** | **0.11** | **0.12** | **0.17** | **0.24** | **0.14** | **0.13** |

**Supplementary Table 12 - FLIM raw data**

### Supplementary Table 13 - Arabidopsis FLIM raw data

| date | measurement # | mVenus fluorescence lifetime [ns] | |
|---|---|---|---|
| | | WOX5-mV | + PLT3-mCh |
| 27.02.2018 | 1 | 3.055 | 3.01 |
| | 2 | 3.076 | 2.93 |
| | 3 | 3.04 | 2.983 |
| | 4 | 3.01 | 3 |
| | 5 | 3.03 | 2.98 |
| | 6 | 3.026 | 2.95 |
| | 7 | 3.06 | 2.96 |
| | 8 | 3.025 | 2.976 |
| | 9 | 3.037 | 2.93 |
| | 10 | 3.007 | 2.941 |
| | 11 | 2.97 | 2.95 |
| | 12 | 3.027 | 2.97 |
| | 13 | 3.025 | 2.96 |
| | 14 | 2.965 | 2.98 |
| | 15 | 2.96 | 2.99 |
| | 16 | 3.011 | 2.991 |
| | 17 | 2.988 | 2.983 |
| | 18 | 2.97 | 2.94 |
| | 19 | 3.01 | 2.97 |
| | 20 | 2.99 | 2.97 |
| | 21 | 2.988 | 2.953 |
| | 22 | 2.964 | 2.92 |
| | 23 | 2.98 | 2.94 |
| | 24 | 2.973 | 2.98 |
| | 25 | | 2.964 |
| 27.04.2018 | 1 | 2.951 | 2.994 |
| | 2 | 2.936 | 2.9 |
| | 3 | 2.98 | 2.7 |
| | 4 | 2.93 | 2.97 |
| | 5 | 2.97 | 2.969 |
| | 6 | 2.987 | 2.951 |
| | 7 | 2.972 | 2.995 |
| | 8 | 2.95 | 2.94 |
| | 9 | 2.97 | 2.92 |
| | 10 | 2.97 | 2.91 |
| | 11 | 2.99 | 2.969 |
| | 12 | 3 | 2.92 |
| | 13 | 2.996 | 2.929 |
| | 14 | 2.97 | 2.9 |
| | 15 | 2.9 | 2.957 |
| | 16 | 2.94 | 2.98 |
| | 17 | 3 | 2.97 |
| | 18 | 2.973 | 2.992 |
| | 19 | 2.95 | 3.001 |
| | 20 | 2.947 | 2.95 |
| | 21 | 2.95 | 2.98 |
| | 22 | | 2.96 |
| | 23 | | 2.91 |

### Supplementary Table 13 - Arabidopsis FLIM raw data

| | | | |
|---|---|---|---|
| | 24 | | 2.9 |
| **25.07.2018** | 1 | 3.2 | 2.88 |
| | 2 | 2.964 | 2.85 |
| | 3 | 2.931 | 2.88 |
| | 4 | 2.984 | 2.9 |
| | 5 | 2.9 | 2.909 |
| | 6 | 2.91 | 2.925 |
| | 7 | 2.965 | 2.9 |
| | 8 | 2.892 | 2.86 |
| | 9 | 2.87 | 2.85 |
| | 10 | 2.8 | |
| | 11 | 2.95 | |
| | 12 | 2.956 | |
| | 13 | 2.933 | |
| | 14 | 2.891 | |
| **27.08.2018** | 1 | 3 | 3.049 |
| | 2 | 2.984 | 3.038 |
| | 3 | 2.72 | 3.024 |
| | 4 | 2.91 | 2.79 |
| | 5 | 2.968 | 2.706 |
| | 6 | 2.978 | 2.67 |
| | 7 | 2.72 | 3 |
| | 8 | 2.92 | 2.99 |
| | 9 | | 2.69 |
| | 10 | | 2.985 |
| **Mean** | | 2.97 | 2.94 |
| **St.dev.** | | 0.07 | 0.08 |

### Supplementary Table 14 - SCN raw data rescue experiments

| | | *Col* | | *plt2* | | *plt3* | | *plt3; pPLT3::PLT3-mV* | | *plt2,3* | | *plt2,3; pPLT3::PLT3-mV* | | *plt2,3; pPLT3::PLT3dPrD-mV* | | *wox5* | | *wox5; pWOX5::WOX5-mV* | |
|---|---|---|---|---|---|---|---|---|---|---|---|---|---|---|---|---|---|---|---|
| **date** | **root #** | QC divisions | CSC layers | QC divisions | CSC layers | QC divisions | CSC layers | QC divisions | CSC layers | QC divisions | CSC layers | QC divisions | CSC layers | QC divisions | CSC layers | QC divisions | CSC layers | QC divisions | CSC layers |
| **2019-08-01** | **1** | 1 | 1 | | | 1 | 1 | 0 | 1 | | | | | | | | | | |
| | **2** | 0 | 0 | | | 2 | 1 | 1 | 1 | | | | | | | | | | |
| | **3** | 0 | 1 | | | 1 | 1 | 2 | 1 | | | | | | | | | | |
| | **4** | 1 | 1 | | | 1 | 1 | 1 | 1 | | | | | | | | | | |
| | **5** | 0 | 1 | | | 4 | 0 | 0 | 1 | | | | | | | | | | |
| | **6** | 0 | 1 | | | 1 | 1 | 0 | 1 | | | | | | | | | | |
| | **7** | 0 | 2 | | | 0 | 0 | 3 | 1 | | | | | | | | | | |
| | **8** | 0 | 1 | | | 1 | 2 | 0 | 1 | | | | | | | | | | |
| | **9** | 0 | 1 | | | 4 | 0 | 1 | 1 | | | | | | | | | | |
| | **10** | 0 | 1 | | | 1 | 1 | 1 | 1 | | | | | | | | | | |
| | **11** | 0 | 1 | | | 3 | 1 | 0 | 0 | | | | | | | | | | |
| | **12** | 2 | 1 | | | 1 | 2 | 0 | 1 | | | | | | | | | | |
| | **13** | 2 | 1 | | | 1 | 1 | 1 | 1 | | | | | | | | | | |
| | **14** | 0 | 2 | | | 0 | 1 | 3 | 0 | | | | | | | | | | |
| | **15** | 1 | 1 | | | 2 | 1 | 1 | 1 | | | | | | | | | | |
| | **16** | 0 | 1 | | | 1 | 1 | 2 | 1 | | | | | | | | | | |
| | **17** | 1 | 1 | | | 1 | 1 | 1 | 1 | | | | | | | | | | |
| | **18** | 0 | 1 | | | 3 | 1 | 0 | 1 | | | | | | | | | | |
| | **19** | 1 | 1 | | | 2 | 0 | 1 | 1 | | | | | | | | | | |
| | **20** | 0 | 1 | | | 2 | 0 | 0 | 1 | | | | | | | | | | |
| | **21** | 0 | 1 | | | | | 0 | 0 | | | | | | | | | | |
| | **22** | 1 | 1 | | | | | 1 | 1 | | | | | | | | | | |
| | **23** | 1 | 1 | | | | | 0 | 1 | | | | | | | | | | |
| | **24** | 0 | 1 | | | | | 1 | 1 | | | | | | | | | | |
| | **25** | 0 | 1 | | | | | 1 | 1 | | | | | | | | | | |
| | **26** | 1 | 0 | | | | | 0 | 0 | | | | | | | | | | |
| | **27** | | | | | | | 0 | 1 | | | | | | | | | | |
| | **28** | | | | | | | 2 | 1 | | | | | | | | | | |
| | **29** | | | | | | | 1 | 0 | | | | | | | | | | |
| | **30** | | | | | | | 1 | 2 | | | | | | | | | | |
| | **31** | | | | | | | 0 | 1 | | | | | | | | | | |
| **2019-05-01** | | QC divisions | CSC layers | QC divisions | CSC layers | QC divisions | CSC layers | QC divisions | CSC layers | QC divisions | CSC layers | QC divisions | CSC layers | QC divisions | CSC layers | QC divisions | CSC layers | QC divisions | CSC layers |
| | **1** | 0 | 1 | | | 1 | 0 | 0 | 1 | 0 | 1 | | | | | 2 | 0 | 2 | 0 |
| | **2** | 0 | 1 | | | 0 | 0 | 0 | 0 | 0 | 1 | | | | | 2 | 0 | 0 | 0 |
| | **3** | 0 | 1 | | | 1 | 0 | 3 | 0 | 1 | 0 | | | | | 4 | 0 | 0 | 1 |
| | **4** | 0 | 0 | | | 0 | 0 | 0 | 1 | 1 | 0 | | | | | 2 | 0 | 0 | 2 |
| | **5** | 0 | 1 | | | 2 | 0 | 0 | 0 | 1 | 3 | | | | | 3 | 0 | 0 | 0 |
| | **6** | 0 | 2 | | | 0 | 0 | 0 | 0 | 0 | 2 | | | | | 4 | 0 | 0 | 0 |
| | **7** | 0 | 1 | | | 0 | 1 | 1 | 1 | 2 | 2 | | | | | 0 | 1 | 0 | 1 |
| | **8** | 0 | 1 | | | 0 | 1 | 1 | 1 | 0 | 1 | | | | | 2 | 0 | 0 | 2 |
| | **9** | 0 | 1 | | | 1 | 0 | 0 | 1 | 4 | 2 | | | | | 1 | 0 | 0 | 2 |
| | **10** | 0 | 1 | | | 1 | 0 | 1 | 0 | 2 | 0 | | | | | 1 | 0 | 0 | 1 |
| | **11** | 0 | 0 | | | 1 | 1 | 0 | 2 | 0 | 0 | | | | | 3 | 1 | 0 | 0 |
| | **12** | 0 | 1 | | | 1 | 0 | 2 | 0 | 0 | 0 | | | | | 1 | 1 | 0 | 0 |
| | **13** | 1 | 1 | | | 1 | 1 | 1 | 0 | 1 | 0 | | | | | 1 | 0 | 0 | 2 |
| | **14** | 0 | 1 | | | 0 | 0 | 0 | 2 | 3 | 0 | | | | | 2 | 0 | 0 | 0 |
| | **15** | 0 | 0 | | | 0 | 0 | 0 | 0 | 2 | 0 | | | | | 3 | 0 | 0 | 1 |
| | **16** | 0 | 1 | | | 1 | 0 | 2 | 0 | 1 | 1 | | | | | 1 | 1 | 0 | 2 |
| | **17** | 0 | 1 | | | 1 | 0 | 0 | 2 | 0 | 0 | | | | | 0 | 1 | 0 | 1 |
| | **18** | 1 | 0 | | | 2 | 0 | 1 | 0 | 1 | 0 | | | | | 1 | 0 | 0 | 0 |
| | **19** | 0 | 2 | | | 0 | 1 | 0 | 1 | 1 | 1 | | | | | 3 | 0 | 0 | 2 |
| | **20** | 0 | 1 | | | 0 | 0 | 3 | 1 | 3 | 2 | | | | | 1 | 2 | 0 | 1 |
| | **21** | 1 | 1 | | | | | 0 | 0 | 1 | 2 | | | | | 1 | 0 | 0 | 1 |
| | **22** | 1 | 0 | | | | | 1 | 1 | 2 | 0 | | | | | 0 | 0 | | |
| | **23** | 1 | 1 | | | | | 1 | 1 | 1 | 0 | | | | | 0 | 1 | | |
| | **24** | 0 | 1 | | | | | 0 | 2 | | | | | | | 2 | 0 | | |
| | **25** | 0 | 0 | | | | | 1 | 0 | | | | | | | 3 | 0 | | |
| | **26** | 2 | 0 | | | | | 1 | 1 | | | | | | | 2 | 0 | | |
| | **27** | 2 | 0 | | | | | 0 | 2 | | | | | | | 2 | 1 | | |
| | **28** | 0 | 1 | | | | | 0 | 0 | | | | | | | 1 | 1 | | |
| | **29** | 0 | 1 | | | | | 0 | 1 | | | | | | | 2 | 0 | | |
| | **30** | 1 | 2 | | | | | 0 | 1 | | | | | | | 3 | 0 | | |
| | **31** | | | | | | | 0 | 0 | | | | | | | | | | |
| | **32** | | | | | | | 1 | 0 | | | | | | | | | | |
| | **33** | | | | | | | 1 | 0 | | | | | | | | | | |
| | **34** | | | | | | | 0 | 1 | | | | | | | | | | |
| | **35** | | | | | | | 2 | 1 | | | | | | | | | | |
| | **36** | | | | | | | 2 | 2 | | | | | | | | | | |
| | **37** | | | | | | | 1 | 1 | | | | | | | | | | |
| | **38** | | | | | | | 1 | 0 | | | | | | | | | | |
| | **39** | | | | | | | 2 | 1 | | | | | | | | | | |
| | **40** | | | | | | | 0 | 2 | | | | | | | | | | |
| | **41** | | | | | | | 0 | 1 | | | | | | | | | | |

### Supplementary Table 14 - SCN raw data rescue experiments

| | | | | | | | | | | | | | | | | | | | |
|---|---|---|---|---|---|---|---|---|---|---|---|---|---|---|---|---|---|---|---|
| | 42 | | | | | | | 2 | 1 | | | | | | | | | | |
| | 43 | | | | | | | 1 | 1 | | | | | | | | | | |
| | 44 | | | | | | | 1 | 0 | | | | | | | | | | |
| | 45 | | | | | | | | | | | | | | | | | | |
| | 46 | | | | | | | | | | | | | | | | | | |
| | 47 | | | | | | | | | | | | | | | | | | |
| | 48 | | | | | | | | | | | | | | | | | | |
| 2018-08-01 | | QC divisions | CSC layers | QC divisions | CSC layers | QC divisions | CSC layers | QC divisions | CSC layers | QC divisions | CSC layers | QC divisions | CSC layers | QC divisions | CSC layers | QC divisions | CSC layers | QC divisions | CSC layers |
| | 1 | | | 1 | 1 | | | 0 | 1 | | | 0 | 1 | 0 | 0 | | | 1 | 1 |
| | 2 | | | 3 | 1 | | | 1 | 1 | | | 1 | 1 | 2 | 0 | | | 0 | 0 |
| | 3 | | | 0 | 1 | | | 2 | 1 | | | 2 | 0 | 2 | 0 | | | 0 | 1 |
| | 4 | | | 1 | 1 | | | 3 | 2 | | | 0 | 1 | 0 | 1 | | | 2 | 1 |
| | 5 | | | 2 | 1 | | | 1 | 1 | | | 2 | 0 | 2 | 0 | | | 0 | 1 |
| | 6 | | | 2 | 0 | | | 2 | 2 | | | 3 | 1 | 2 | 0 | | | 1 | 1 |
| | 7 | | | 1 | 0 | | | 2 | 1 | | | 2 | 1 | 0 | 1 | | | 1 | 1 |
| | 8 | | | 2 | 1 | | | 1 | 2 | | | 2 | 1 | 2 | 0 | | | 2 | 1 |
| | 9 | | | 1 | 0 | | | 1 | 1 | | | 0 | 1 | 0 | 1 | | | 0 | 1 |
| | 10 | | | 1 | 2 | | | 0 | 1 | | | 1 | 1 | 1 | 1 | | | 0 | 1 |
| | 11 | | | 1 | 1 | | | 1 | 1 | | | 4 | 1 | 1 | 1 | | | 2 | 0 |
| | 12 | | | 1 | 1 | | | 0 | 1 | | | 2 | 2 | 2 | 0 | | | 1 | 0 |
| | 13 | | | 2 | 1 | | | 4 | 0 | | | 2 | 0 | 3 | 0 | | | 1 | 1 |
| | 14 | | | 3 | 0 | | | 2 | 2 | | | 0 | 0 | 1 | 0 | | | 0 | 1 |
| | 15 | | | 0 | 2 | | | 0 | 1 | | | 0 | 1 | 1 | 0 | | | 0 | 1 |
| | 16 | | | 1 | 2 | | | 3 | 1 | | | 2 | 1 | | | | | 1 | 1 |
| | 17 | | | 1 | 1 | | | 0 | 1 | | | 3 | 0 | | | | | 1 | 1 |
| | 18 | | | 1 | 1 | | | 1 | 1 | | | 0 | 0 | | | | | 2 | 0 |
| | 19 | | | 2 | 0 | | | 1 | 2 | | | 1 | 0 | | | | | 1 | 2 |
| | 20 | | | 1 | 0 | | | 3 | 1 | | | 1 | 0 | | | | | 1 | 2 |
| | 21 | | | 0 | 0 | | | 0 | 1 | | | 1 | 1 | | | | | 0 | 1 |
| | 22 | | | | | | | 1 | 1 | | | 0 | 0 | | | | | 0 | 1 |
| | 23 | | | | | | | 3 | 0 | | | 3 | 2 | | | | | 1 | 0 |
| | 24 | | | | | | | 0 | 1 | | | 3 | 0 | | | | | 0 | 1 |
| | 25 | | | | | | | 2 | 1 | | | 3 | 0 | | | | | 1 | 0 |
| | 26 | | | | | | | 0 | 2 | | | 1 | 0 | | | | | 1 | 1 |
| | 27 | | | | | | | 1 | 0 | | | 0 | 1 | | | | | 0 | 1 |
| | 28 | | | | | | | 4 | 0 | | | 2 | 1 | | | | | 0 | 1 |
| | 29 | | | | | | | 2 | 1 | | | 0 | 0 | | | | | 0 | 1 |
| | 30 | | | | | | | | | | | 2 | 0 | | | | | 0 | 1 |
| 2021-07-07 | | QC divisions | CSC layers | QC divisions | CSC layers | QC divisions | CSC layers | QC divisions | CSC layers | QC divisions | CSC layers | QC divisions | CSC layers | QC divisions | CSC layers | QC divisions | CSC layers | QC divisions | CSC layers |
| | 1 | 1 | 1 | 0 | 1 | | | | | 0 | 0 | 2 | 1 | 1 | 2 | | | | |
| | 2 | 2 | 1 | 2 | 2 | | | | | 3 | 1 | 0 | 0 | 1 | 1 | | | | |
| | 3 | 2 | 0 | 0 | 1 | | | | | 1 | 1 | 3 | 1 | 1 | 1 | | | | |
| | 4 | 1 | 0 | 2 | 1 | | | | | 2 | 0 | 2 | 1 | 1 | 1 | | | | |
| | 5 | 3 | 1 | 2 | 1 | | | | | 0 | 0 | 1 | 1 | 3 | 0 | | | | |
| | 6 | 1 | 1 | 1 | 1 | | | | | 1 | 0 | 1 | 1 | 1 | 0 | | | | |
| | 7 | 0 | 1 | 3 | 1 | | | | | 1 | 0 | 1 | 1 | 2 | 0 | | | | |
| | 8 | 2 | 1 | 2 | 1 | | | | | 3 | 1 | 2 | 1 | 3 | 0 | | | | |
| | 9 | 0 | 2 | 2 | 1 | | | | | 3 | 1 | 2 | 1 | 1 | 0 | | | | |
| | 10 | 0 | 1 | | | | | | | 1 | 1 | 3 | 0 | 4 | 0 | | | | |
| | 11 | 0 | 1 | | | | | | | 0 | 1 | 1 | 0 | 2 | 1 | | | | |
| | 12 | 2 | 1 | | | | | | | 2 | 0 | 0 | 1 | 3 | 0 | | | | |
| | 13 | 1 | 0 | | | | | | | 0 | 0 | 1 | 1 | 3 | 0 | | | | |
| | 14 | 0 | 1 | | | | | | | 1 | 1 | 2 | 2 | 1 | 1 | | | | |
| | 15 | 0 | 0 | | | | | | | 2 | 0 | 2 | 1 | | | | | | |
| | 16 | 1 | 1 | | | | | | | 3 | 2 | | | | | | | | |
| | 17 | 1 | 2 | | | | | | | 2 | 0 | | | | | | | | |
| | 18 | 0 | 0 | | | | | | | 1 | 0 | | | | | | | | |
| | 19 | 1 | 0 | | | | | | | 2 | 0 | | | | | | | | |
| | 20 | 3 | 2 | | | | | | | 2 | 0 | | | | | | | | |
| | 21 | 1 | 2 | | | | | | | 2 | 0 | | | | | | | | |
| | 22 | 2 | 2 | | | | | | | 1 | 1 | | | | | | | | |
| | 23 | 0 | 2 | | | | | | | 1 | 0 | | | | | | | | |
| | 24 | 0 | 1 | | | | | | | | | | | | | | | | |
| | 25 | 3 | 0 | | | | | | | | | | | | | | | | |
| | 26 | 1 | 1 | | | | | | | | | | | | | | | | |
| | 27 | 1 | 0 | | | | | | | | | | | | | | | | |
| | 28 | 0 | 0 | | | | | | | | | | | | | | | | |
| | 29 | 1 | 1 | | | | | | | | | | | | | | | | |
| | 30 | 1 | 1 | | | | | | | | | | | | | | | | |
| 2021-07-27 | | QC divisions | CSC layers | QC divisions | CSC layers | QC divisions | CSC layers | QC divisions | CSC layers | QC divisions | CSC layers | QC divisions | CSC layers | QC divisions | CSC layers | QC divisions | CSC layers | QC divisions | CSC layers |
| | 1 | 1 | 0 | 0 | 0 | | | | | 0 | 0 | 2 | 2 | 2 | 0 | | | | |
| | 2 | 2 | 2 | 0 | 1 | | | | | 0 | 1 | 1 | 0 | 0 | 1 | | | | |
| | 3 | 0 | 0 | 0 | 0 | | | | | 1 | 0 | 1 | 0 | 0 | 1 | | | | |
| | 4 | 0 | 1 | 0 | 1 | | | | | 3 | 2 | 2 | 0 | 1 | 1 | | | | |
| | 5 | 2 | 1 | 0 | 2 | | | | | 0 | 0 | 0 | 0 | 2 | 0 | | | | |

### Supplementary Table 14 - SCN raw data rescue experiments

| | | | | | | | | | | | | | | | | | | |
|---|---|---|---|---|---|---|---|---|---|---|---|---|---|---|---|---|---|---|
| 6 | 0 | 1 | 1 | 0 | | | | | 1 | 0 | 3 | 1 | 0 | 2 | | | | |
| 7 | 1 | 1 | 0 | 0 | | | | | 2 | 1 | 0 | 0 | 0 | 0 | | | | |
| 8 | 0 | 1 | 0 | 1 | | | | | 1 | 1 | 2 | 2 | 3 | 0 | | | | |
| 9 | 0 | 0 | 0 | 1 | | | | | 3 | 0 | 1 | 0 | 2 | 1 | | | | |
| 10 | 0 | 1 | | | | | | | 1 | 0 | 1 | 1 | 1 | 1 | | | | |
| 11 | 1 | 1 | | | | | | | 2 | 1 | 3 | 1 | 0 | 0 | | | | |
| 12 | 1 | 0 | | | | | | | 1 | 0 | 0 | 1 | 0 | 0 | | | | |
| 13 | 0 | 2 | | | | | | | 2 | 0 | 0 | 0 | 0 | 0 | | | | |
| 14 | 0 | 0 | | | | | | | 0 | 0 | 1 | 1 | 2 | 2 | | | | |
| 15 | 0 | 2 | | | | | | | 0 | 1 | 1 | 0 | 0 | 0 | | | | |
| 16 | 0 | 1 | | | | | | | 0 | 0 | 1 | 1 | | | | | | |
| 17 | 1 | 1 | | | | | | | 0 | 1 | 2 | 0 | | | | | | |
| 18 | 0 | 0 | | | | | | | 3 | 0 | | | | | | | | |
| 19 | 0 | 0 | | | | | | | 1 | 0 | | | | | | | | |
| 20 | 0 | 1 | | | | | | | | | | | | | | | | |
| **mean** | **0.58** | **0.89** | **1.08** | **0.85** | **1.13** | **0.55** | **0.95** | **0.88** | **1.26** | **0.55** | **1.40** | **0.66** | **1.34** | **0.48** | **1.77** | **0.33** | **0.43** | **0.88** |
| **SD** | **0.79** | **0.60** | **0.94** | **0.62** | **1.03** | **0.59** | **1.01** | **0.62** | **1.06** | **0.74** | **1.05** | **0.62** | **1.09** | **0.62** | **1.12** | **0.54** | **0.66** | **0.65** |
| **n** | **106** | **106** | **39** | **39** | **40** | **40** | **104** | **104** | **65** | **65** | **62** | **62** | **44** | **44** | **30** | **30** | **51** | **51** |
